# Single cell autofluorescence imaging reveals immediate metabolic shifts of neutrophils with activation across biological systems

**DOI:** 10.1101/2024.07.26.605362

**Authors:** Rupsa Datta, Veronika Miskolci, Gina M. Gallego-López, Emily Britt, Amani Gillette, Aleksandr Kralovec, Morgan A. Giese, Tongcheng Qian, James Votava, Jing Fan, Anna Huttenlocher, Melissa Skala

## Abstract

Neutrophils, the most abundant leukocytes in human peripheral circulation, are crucial for the innate immune response. They are typically quiescent but rapidly activate in response to infection and inflammation, performing diverse functions such as oxidative burst, phagocytosis, and NETosis, which require significant metabolic adaptation. Deeper insights into such metabolic changes will help identify regulation of neutrophil functions in health and diseases. Due to their short lifespan and associated technical challenges, the metabolic processes of neutrophils are not completely understood. This study uses optical metabolic imaging (OMI), which entails optical redox ratio and fluorescence lifetime imaging microscopy of intrinsic metabolic coenzymes NAD(P)H and FAD to assess the metabolic state of single neutrophils. Primary human neutrophils were imaged *in vitro* under a variety of activation conditions and metabolic pathway inhibitors, while metabolic and functional changes were confirmed with mass spectrometry, oxidative burst, and NETosis measurements. Our findings show that neutrophils undergo rapid metabolic remodeling to a reduced redox state indicated by changes in NAD(P)H lifetime and optical redox ratio, with a shift to an oxidized redox state during activation. Additionally, single cell OMI analysis reveals a heterogeneous metabolic response across neutrophils and human donors to live pathogen infection (*Pseudomonas aeruginosa* and *Toxoplasma gondii*). Finally, consistent OMI changes with activation were confirmed between *in vitro* human and *in vivo* zebrafish larvae neutrophils. This study demonstrates the potential of OMI as a versatile tool for studying neutrophil metabolism and underscores its use across different biological systems, offering insights into neutrophil metabolic activity and function at a single cell level.

## Introduction

Neutrophils, also known as polymorphonuclear cells, are the most abundant leukocytes in human peripheral circulation. They are part of the innate immune system and are the first to arrive at sites of infection and inflammation. Neutrophils in circulation are generally quiescent, and their activation is an essential component of a robust innate immune system. In response to inflammatory cues and infections, neutrophils undergo rapid differentiation through granulopoiesis in the bone marrow, followed by the mobilization of neutrophils to the infection site. Upon activation, neutrophils combat infiltrating pathogens through a variety of effector functions, including a production of reactive oxygen species (ROS) known as oxidative burst, phagocytosis, and the formation and expulsion of neutrophil extracellular traps (NETs), or NETosis^1^.

These processes are associated with increased energy demand. Additionally, metabolic preferences have been linked to neutrophil inflammatory potential, apoptosis, biosynthesis, maturation, and epigenetics, underscoring the significance of cellular metabolism in neutrophil function^2^. However, unlike other immune cells, knowledge of neutrophil metabolism is still at its nascency^3–5^. Previous studies have predominantly emphasized glycolysis as the primary metabolic pathway for deriving energy in neutrophils, leaving the involvement of other metabolic pathways in neutrophil activation and function less explored^2,3^. Recent advances indicate that the high metabolic demand of activation also requires remodeling of other metabolic pathways including the pentose phosphate pathway (PPP) or the hexose monophosphate shunt, tricarboxylic acid (TCA) cycle, fatty acid oxidation, and oxidative phosphorylation^3,5^. Despite having a short lifespan, these terminally differentiated cells exhibit metabolic flexibility across diverse environments and require fast metabolic remodeling, especially during effector functions^4,6^.

Aberrant metabolism in neutrophils holds substantial importance in various disease contexts. For example, glucose-6-phosphate dehydrogenase (G6PD) deficiency is characterized by reduced NADPH production that results in effector function defects^3^. Additionally, dysfunctional neutrophil metabolism could either result in reduced functions like NETosis^7^ (e.g., severe COVID-19) or excessive ROS generation, cytotoxic protease release, and excessive NETosis that ultimately lead to tissue damage (e.g., rheumatoid arthritis and systemic lupus erythematosus)^3^. Given the increasing relevance of understanding neutrophil metabolism in disease contexts, there is a growing need to obtain metabolic information. Moreover, previous research suggests that single-cell heterogeneity is a critical yet poorly understood feature of neutrophil function, underscoring the need for metabolic analysis at the single-cell level^8–10^. Finally, intracellular reactions related to neutrophil activation occur at a rapid rate necessitating fast readout techniques ^4,11^. Current techniques to study neutrophil metabolism such as mass spectrometry, enzyme activity measurements, and extracellular flux analysis^7,12^ are destructive and measure the activity of the bulk population. This emphasizes the need for non-invasive single-cell techniques to measure metabolism in neutrophils, both *in vitro* and *in vivo*. Additionally, there is a lack of techniques that can capture real-time, rapid changes in neutrophil metabolism. Finally, no technique currently exists that can accurately recognize and classify neutrophil activation based on metabolic biomarkers.

Our work addresses these needs with optical metabolic imaging (OMI), a label-free technique capable of monitoring rapid metabolic changes at the single-cell level in neutrophils. OMI entails multiphoton microscopy of intrinsic fluorophores, reduced nicotinamide adenine dinucleotide (phosphate) (NAD(P)H) and oxidized flavin adenine dinucleotide (FAD). The non-invasive nature of this approach enables rapid repeated measurements of metabolic dynamics in single cells. NAD(P)H and FAD participate directly in oxidation-reduction reactions in metabolic pathways and can be used to monitor the metabolic status of cells. We measure the optical redox ratio, defined as the fluorescence intensity of NAD(P)H/(NAD(P)H+FAD), which describes the redox state of cells^13^. Additionally, we perform fluorescence lifetime imaging microscopy (FLIM), which provides information on the enzyme binding status of the fluorophores^14^.

While OMI has been previously used to study the electron transport chain in mitochondrial respiration^15^, this work also uses OMI to investigate electron transport across membranes by the NADPH oxidase (NOX)2 enzymatic complex^16,17^. In activated neutrophils, the membrane protein NOX2 (or gp91phox) – the main subunit of the NOX2 complex – plays an essential role in oxidative burst and other effector functions. It is the primary production site of reactive oxygen species (ROS), including superoxide radical anion (O_2_ ^-^), and subsequently hydrogen peroxide (H_2_O_2_) and hypochlorous acid (HOCl)^17,18^. To generate superoxide, the NOX2 system functions as a transmembrane electron transfer chain where electron flow occurs from NADPH via FAD, inner and outer membrane heme groups, and finally to molecular oxygen in seven steps^16^. The C-terminal of NOX2 provides binding sites for both NADPH and FAD that are essential for electron flow in the oxidase^16,17^.

In this study we use single-cell OMI (optical redox ratio and FLIM of NAD(P)H and FAD) to show that neutrophils remodel their metabolism within minutes upon activation. Specifically, we find a rapid transition to reduced redox state that gradually shifts towards an oxidized state, confirmed with metabolite analysis, after the onset of the oxidative burst and NETosis. Additionally, single-cell metabolic analysis revealed heterogenous response of neutrophils to live pathogens. Finally, our study highlights the consistency of *in vitro* and *in vivo* neutrophil responses to activation. We demonstrate that OMI is sensitive to metabolic remodeling in activated primary human neutrophils *in vitro* and in zebrafish larvae *in vivo*. This suggests that OMI can reliably capture metabolic changes across different biological systems. This cross-species applicability enhances the generalizability of our findings and underscores the robustness of OMI as a versatile tool for monitoring single-cell changes over time in neutrophil metabolism.

## Results

### Optical metabolic imaging is sensitive to immediate metabolic shifts in neutrophils upon activation

To confirm the activation of primary human neutrophils (Supp Fig 1a), we analyzed oxidative burst and NETosis in neutrophils treated with the pharmacological stimuli phorbol myristate acetate (PMA) – a protein kinase C activator commonly used as a pro-inflammatory activation agent^19^. PMA treatment did not induce significant neutrophil death by 60min (Supp Fig 1b). The onset of both effector functions was about 30 to 60 minutes after PMA activation (Fig. 1 a-b, Supp Fig 1c. The oxidative burst was not as noticeably induced by other stimuli, such as lipopolysaccharide (LPS), an endotoxin from gram-negative bacteria, and tumor necrosis factor- α (TNF-α), a cytokine, even after a measurement period of 5 hours (Supp Fig 1c). This is consistent with a previous observation in primary human neutrophils^12^. However, some increase in NETosis was observed with TNF-α after 6 hours (Supp Fig.1d).

**Figure 1.**
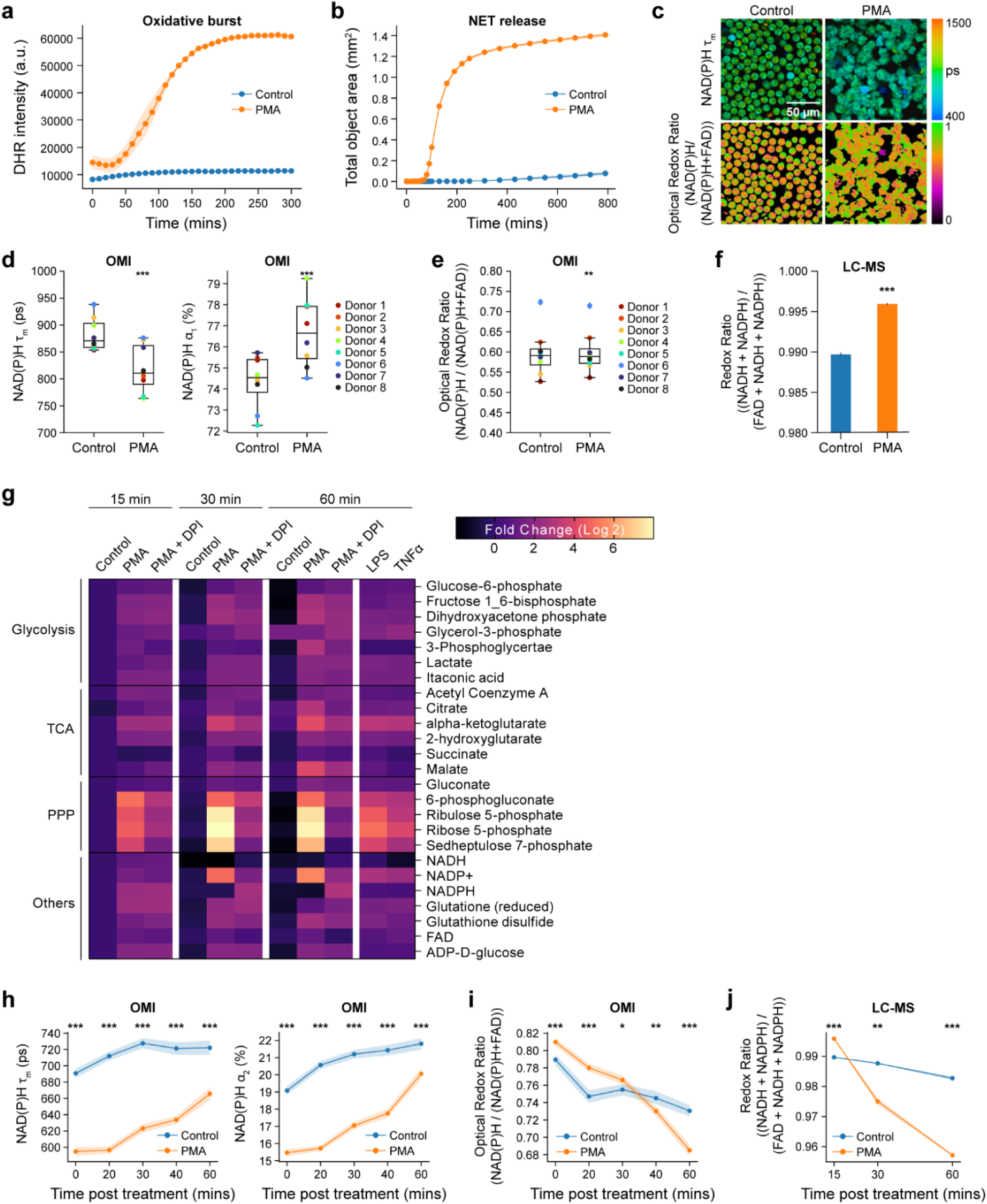
Primary human neutrophils undergo metabolic alteration immediately upon activation. (a) Quantification of fluorescence intensity of Dihydrorhodamine 123 (DHR intensity) indicating intracellular ROS in unstimulated control and PMA treated neutrophils (Donor 10). At least 5 images were acquired per condition every 5 minutes for 5 hours, in triplicates. (b) NET release induced by PMA (Donor 14). (c) Representative images of NAD(P)H mean lifetime and optical redox ratio [I_NAD(P)H_/(I_NAD(P)H_+I_FAD_)] in control and PMA treated neutrophils. (d) NAD(P)H mean lifetime (left), free NAD(P)H α_1_ percentage (right) and (e) optical redox ratio of control and PMA (100nM, 15 minutes) treated neutrophils from 8 distinct donors (Donor 1-8). 5-6 images were acquired per condition, each dot represents the average across all cells per donor (n=number of cells across all donors, Table 1) (f) Redox ratio quantified from molar concentration measured by LC-MS from 2 technical replicates (Donor 8) of control and PMA (100nM, 15 minutes) treated neutrophils. (g) Heat map representation of metabolomic variations across the unstimulated control, PMA treatment (100nM) and PMA together with NOX2 inhibitor DPI (10µM) treatment (PMS+DPI) (Donor 8). Heatmap also includes stimulation with LPS (20µg/L) and TNFα (5µg/L). To align with the OMI conditions, metabolites were extracted at 15, 30 and 60 mins for all the conditions except for LPS and TNFα, which were extracted at 60 mins. Each metabolite abundance is normalized to the control abundance at 15 mins then log base 2 transformed (Log2 (measured Abundance/Control Abundance)) for each experimental condition. Metabolites have been grouped based on metabolic pathways. Single-cell quantification of (h) NAD(P)H mean lifetime (left), bound NAD(P)H α_2_percentage (right) and (i) optical redox ratio of control and PMA treated neutrophils acquired every 15 minutes from 0 minutes (after addition of PMA) to 60 minutes after treatment. Each point in (h-i) represents the average for all cells at the indicated timepoint and error bars represent the 95% confidence interval. At least 5 images were acquired per condition, number of cells per time point in Table 2 (Donor 10). (j) Redox ratio quantified from molar concentration measured by LC-MS, of control and PMA treated neutrophils (Donor 8). Significance was determined for (d-f) using one-way analysis of variance (ANOVA) with post hoc Dunnett’s test and (h-j) using Student’s T test. (\*\*\**P* < 0.001; **P<0.01; \**P* < 0.05).

**Table 1.** Cells analyzed per donor per condition

| Donor # | Condition | n = cells analyzed | Donor # | Condition | n = cells |
| --- | --- | --- | --- | --- | --- |
| Donor 1 | Control | 402 | Donor 6 | Control | 628 |
|  | PMA | 938 |  | PMA | 649 |
|  | LPS | 562 |  | LPS | 577 |
| | TNFα | 635 | | TNFα | 842 |
| Donor 2 | Control | 429 |  | PMA + 2DG | 567 |
|  | PMA | 538 |  | PMA + 6AN | 341 |
|  | LPS | 240 |  | PMA + DPI | 391 |
| | TNFα | 320 | | PMA + NaCN | 383 |
| Donor 3 | Control | 985 |  | PMA + AA | 407 |
|  | PMA | 801 |  | PMA + IAA | 441 |
|  | LPS | 679 | Donor 7 | Control | 429 |
| | TNFα | 1128 | | PMA | 457 |
|  | PMA + 2DG | 1065 | Donor 8 | Control | 515 |
|  | PMA + 6AN | 907 |  | PMA | 888 |
|  | PMA + DPI | 999 |  | LPS | 458 |
| | PMA + NaCN | 938 | | TNFα | 810 |
|  | PMA + AA | 739 |  | PMA + 6AN | 595 |
|  | PMA + IAA | 736 |  | PMA + DPI | 530 |
| Donor 4 | Control | 557 | Donor M | Control | 343 |
|  | PMA | 437 |  | ME49 | 331 |
|  | PMA + 2DG | 282 |  | PAO | 370 |
|  | PMA + 6AN | 560 |  | RH | 445 |
|  | PMA + DPI | 590 | Donor N | Control | 282 |
|  | PMA + NaCN | 481 |  | ME49 | 179 |
|  | PMA + AA | 624 |  | PAO | 357 |
|  | PMA + IAA | 551 |  | RH | 122 |
| Donor 5 | Control | 434 | Donor P | Control | 355 |
|  | PMA | 312 |  | ME49 | 247 |
|  | PMA + 2DG | 688 |  | PAO | 305 |
|  | PMA + 6AN | 794 |  | RH | 323 |
|  | PMA + DPI | 653 |  |  |  |
|  | PMA + NaCN | 927 |  |  |  |
|  | PMA + AA | 615 |  |  |  |
|  | PMA + IAA | 595 |  |  |  |

**Table 2.**
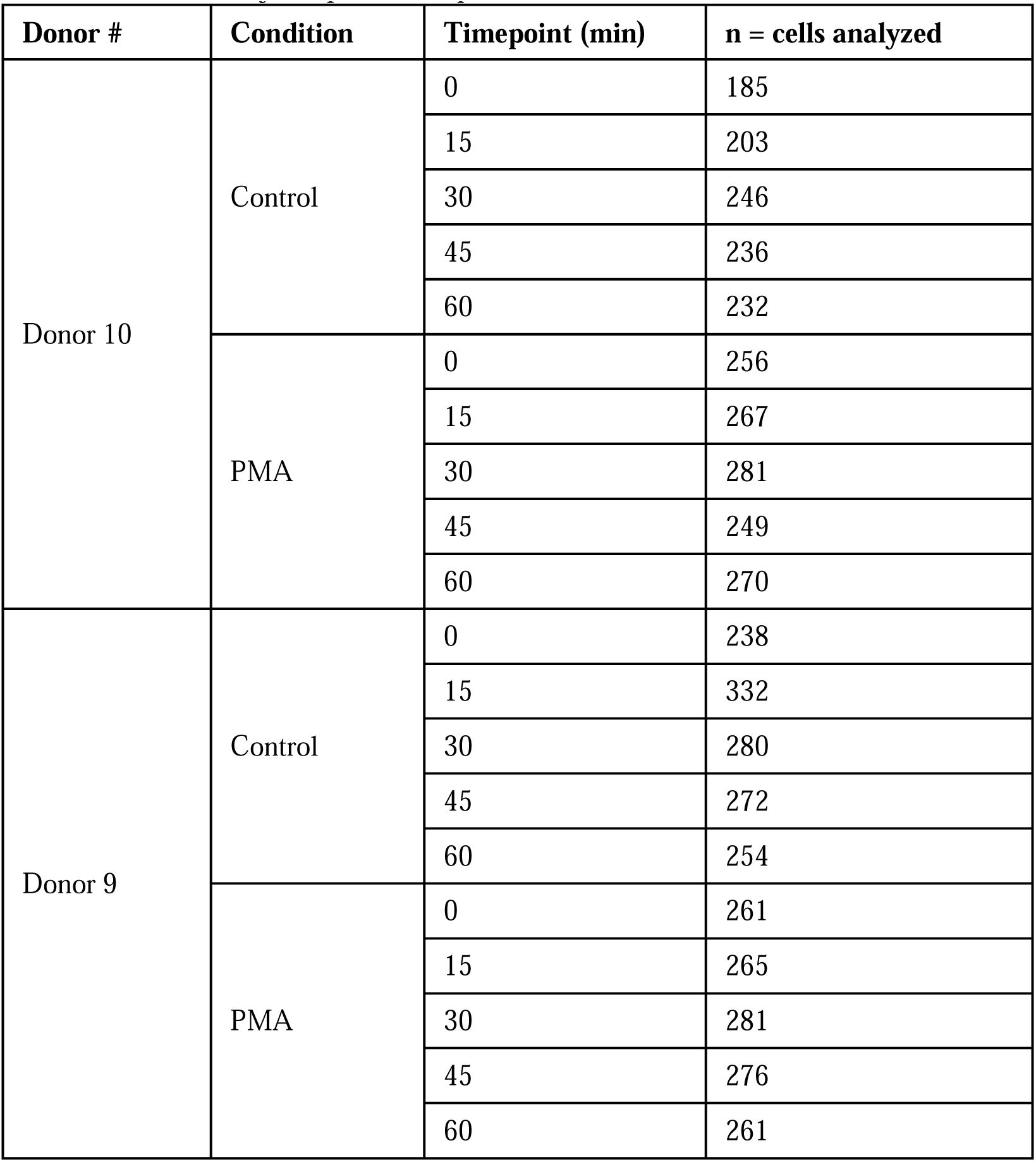
Cells analyzed per donor per condition in timeseries OMI

| Donor # | Condition | Timepoint (min) | n = cells analyzed |
| --- | --- | --- | --- |
| Donor 10 | Control | 0 | 185 |
|  |  | 15 | 203 |
|  |  | 30 | 246 |
|  |  | 45 | 236 |
|  |  | 60 | 232 |
|  | PMA | 0 | 256 |
|  |  | 15 | 267 |
|  |  | 30 | 281 |
|  |  | 45 | 249 |
|  |  | 60 | 270 |
| Donor 9 | Control | 0 | 238 |
|  |  | 15 | 332 |
|  |  | 30 | 280 |
|  |  | 45 | 272 |
|  |  | 60 | 254 |
|  | PMA | 0 | 261 |
|  |  | 15 | 265 |
|  |  | 30 | 281 |
|  |  | 45 | 276 |
|  |  | 60 | 261 |

Next, we sought to investigate the metabolic response of primary human neutrophils to activation using OMI, which leverages the autofluorescence properties of metabolic coenzymes reduced NADH, NADPH, and oxidized FAD. We observed a significant decrease in NAD(P)H mean lifetime and increase in the relative proportion of free NAD(P)H (α_1_ %) within 15 minutes of PMA treatment across all eight donors (Fig 1c-d, Supp Fig 2a, Table 1). In addition, activated neutrophils exhibited a transition towards a reduced redox state (increase in optical redox ratio) in comparison to unstimulated control, consistent across five out of eight donors (Fig 1e, Supp Fig 2b, Supp Fig 5e, Supp Fig 7a, Table 1). The redox ratio calculated from liquid chromatography-mass spectrometry (LC-MS) confirmed this reduced redox state with neutrophil activation in one representative donor (Fig 1f). Despite TNF-α and LPS failing to induce a measurable oxidative burst or NETosis, the activators did elicit a reduction in NAD(P)H mean lifetime in three out of four donors (Supp Fig 2c, 2d, 2f, Table 1). However, changes in the optical redox ratio were not significant with either TNF-α or LPS (Supp Fig 2c, 2e, 2g-h, Table 1).

**Figure 2.**
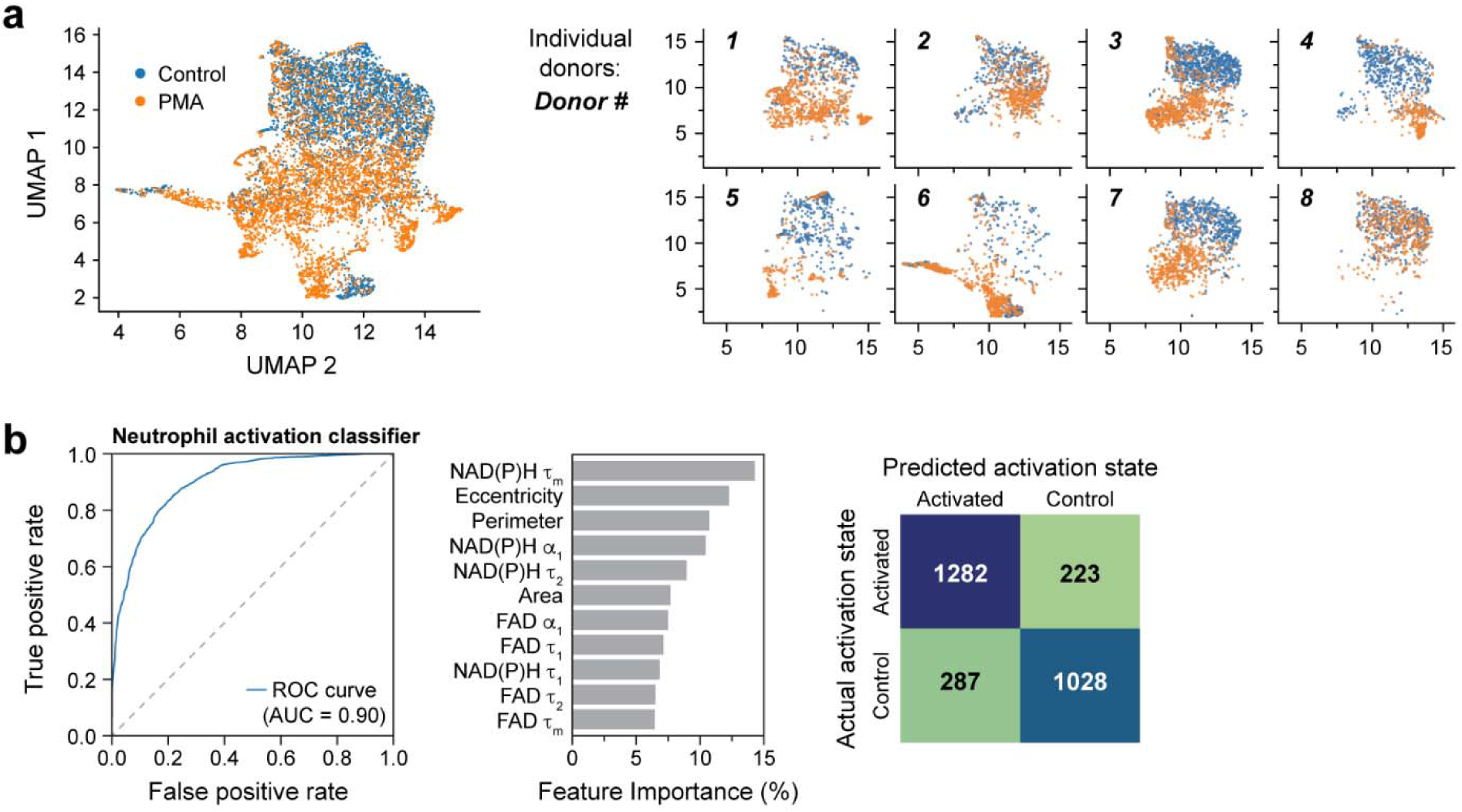
OMI features accurately classify unstimulated and activated primary human neutrophils 15 minutes post-stimulation at the single cell level. (a) UMAP projection based on Euclidean distance metric of parameters (Table 4) of control and PMA treated (100nM, 15 minutes) neutrophils from 8 distinct donors (Donor 1-8). Subplots show datapoints from individual donors. (b) Receiver operating characteristic (ROC) curve (left) and area under the curve (AUC) of random forest model to classify neutrophil activation state and corresponding feature importance (middle) and confusion matrix with number of cells in each category (right). Dish condition was used as ground truth. The classifier was trained and tested on data from donors 1-8, with a split ratio of train: 70%, test: 30% of cells (number of cells in Table 1)

**Table 4.** Variables included in UMAP and classifier

| Figure | Variables |
| --- | --- |
| UMAP (Fig 1j) | NAD(P)H $\alpha 1$,<br>NAD(P)H $\tau 1$,<br>NAD(P)H $\tau 2$,<br>NAD(P)H $\tau m$,<br>FAD $\alpha 1$,<br>FAD $\tau 1$,<br>FAD $\tau 2$,<br>FAD $\tau m$,<br>area,<br>perimeter<br>eccentricity |
| Random forest Classifier (Fig2b) and<br>Coefficient of variation heatmap (Fig 4e) | NAD(P)H $\alpha 1$,<br>NAD(P)H $\tau 1$,<br>NAD(P)H $\tau 2$,<br>NAD(P)H $\tau m$,<br>FAD $\alpha 1$,<br>FAD $\tau 1$,<br>FAD $\tau 2$,<br>FAD $\tau m$,<br>area,<br>perimeter<br>eccentricity |
| Random forest Classifier (Supp Fig 4d) and<br>Coefficient of variation heatmap (Fig 9) | NAD(P)H $\alpha 1$,<br>NAD(P)H $\tau 1$,<br>NAD(P)H $\tau 2$,<br>NAD(P)H $\tau m$,<br>FAD $\alpha 1$,<br>FAD $\tau 1$,<br>FAD $\tau 2$,<br>FAD $\tau m$, |

Taken together, these data suggest an immediate increase in free NAD(P)H upon activation, potentially related to enhanced glycolysis or PPP activity^4^. To confirm this, we analyzed metabolite abundance at 15mins after PMA treatment with LC-MS. Indeed, we found that compared to unstimulated control, PMA-stimulated neutrophils exhibited greater abundance of several glycolytic intermediates (Fig 1g, Supp Fig 3a-b, Table 3). Similarly, we found a higher prevalence of several metabolites from both the oxidative and non-oxidative branches of the PPP within 15 minutes after PMA treatment (Fig 1g) consistent with previous observations^4^. LC-MS results also revealed a slight increase in the abundance of glycolytic and PPP intermediates with TNF-α or LPS activation at 60 minutes (Fig 1g).

**Figure 3.**
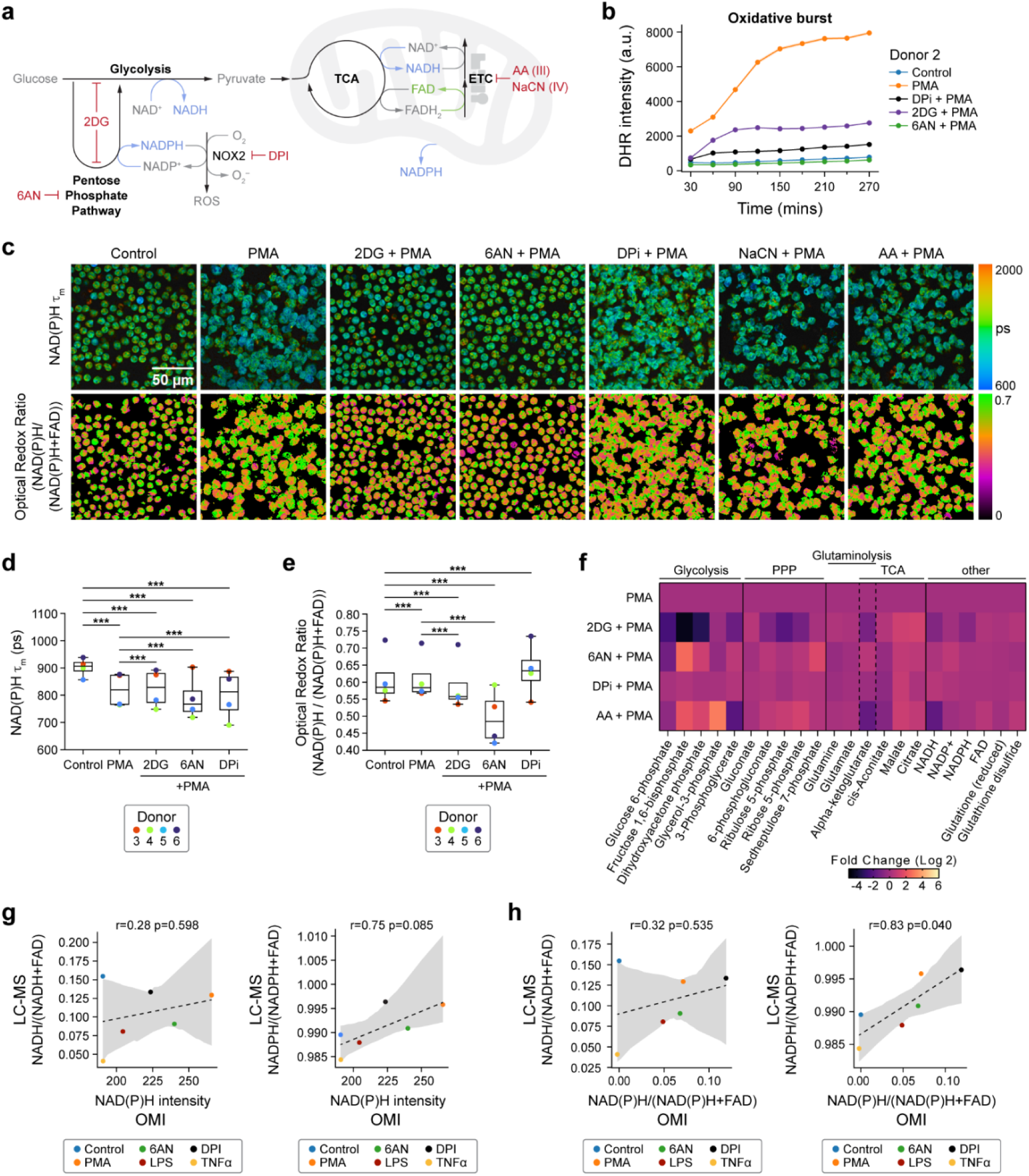
NAD(P)H mean lifetime and optical redox are sensitive to glycolysis and PPP inhibition in activated primary human neutrophils. (a) Illustration of neutrophil metabolic pathways and metabolic inhibitors used. (b) Quantification of fluorescence intensity of Dihydrorhodamine 123 (DHR) indicating intracellular ROS in unstimulated control, PMA- treated, PMA along with inhibitor (2-DG, 6-AN, DPI)-treated neutrophils from one donor (Donor 2). (c) Representative images of NAD(P)H mean lifetime and optical redox ratio. (d) NAD(P)H mean lifetime and (e) optical redox ratio of neutrophils from 4 distinct donors. At least 5 images were acquired per condition, each dot represents the average across all cells per donor (n=number of cells across all donors, Table 1). Significance was determined using one- way analysis of variance (ANOVA) with post hoc Tukey test (\*\*\**P* < 0.001). (f) Heat map representation of metabolomic variations across PMA (100nM) treatment and PMA together with 100mM 2DG (PMS+2DG), 5mM 6AN (PMA+6AN), 10µM DPI (PMA+DPI), and 1µM AA (PMA+AA) (Donor 13). To align with the OMI conditions, metabolites were extracted at 15 minutes. Each metabolite abundance is normalized to the control abundance then log base 2 transformed (Log_2_ (measured Abundance/Control Abundance)) for each experimental condition. Metabolites have been grouped based on metabolic pathways. The results of the significant tests are presented in Table 5. (g-h) Redox ratio quantified from LC-MS measurements (converted to molar concentration) with correlation between LC-MS and OMI measurements performed on neutrophils from one donor (Donor 8). For OMI, 5-6 images were acquired per condition, number of cells in Table 1. (c-h) All data collected at 15 minutes after treatment.

Taking advantage of the fact that label-free OMI allows temporal metabolic measurements of single cells in of the same dish, we tracked neutrophil metabolism from activation to the initiation of effector function (60 minutes after activation with PMA). As observed previously, NAD(P)H mean lifetime was lower immediately upon PMA treatment (Fig 1h, Supp Fig 4a, Table 2). This difference persisted for the measurement duration of 60 minutes. However, the NAD(P)H mean lifetime appeared to increase over time (Fig 1h, Supp Fig. 4a), potentially indicating NAD(P)H binding to NOX2 for subsequent electron transfer to generate ROS, as the enzyme complex starts to form as a result of activation^20^ (Supp Fig 4b). Consequently, we observed an increase in the relative proportion of bound NAD(P)H (α_2_ %) over time (Fig 1h, Supp Fig. 4a). Notably, the initial increase in the optical redox ratio with PMA activation immediately after treatment declined over time, reaching the redox levels of unstimulated cells within 20 to 30 minutes, and cells continued to oxidize thereafter (Fig 1i, Supp Fig 4c).

**Figure 4.**
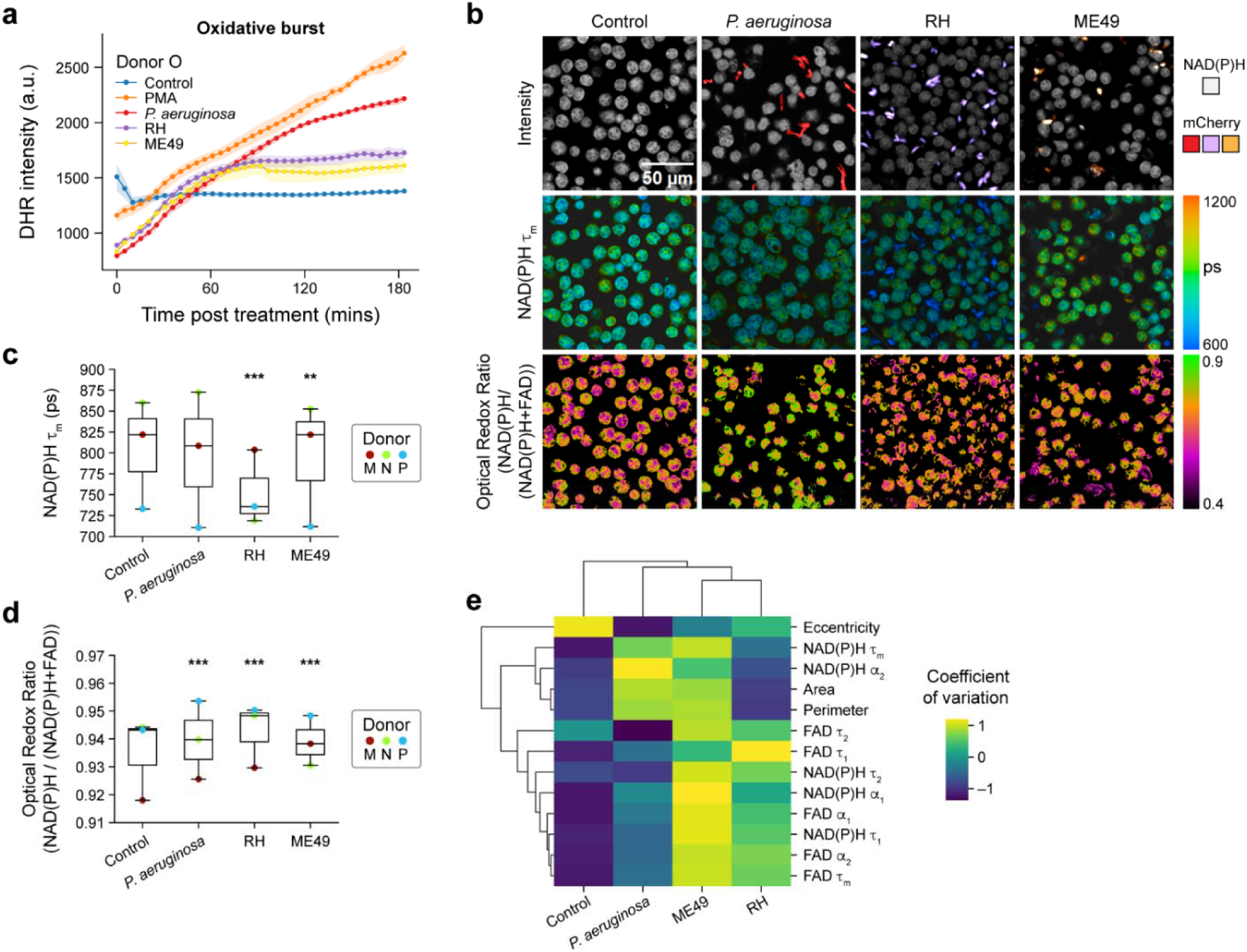
Pathogens elicit immediate and heterogenous metabolic response in primary human neutrophils. (a) Quantification of fluorescence intensity of Dihydrorhodamine 123 (DHR) indicating intracellular ROS in unstimulated control, PMA, coculture with *P. aeruginosa*, and *T. gondii* strains RH and M49 (Donor O). (b) Representative images. (c) NAD(P)H mean lifetime and (d) optical redox ratio of unstimulated control, coculture with *P. aeruginosa*, and *T. gondii* strains RH and ME49 for 15 minutes in neutrophils from 3 distinct donors (Donor M, N and P). At least 5 images were acquired per condition, each dot represents the average across all cells per donor (n=number of cells across all donors, Table 1). Significance was determined using one-way analysis of variance (ANOVA) with post hoc Tukey test (\*\*\**P* < 0.001; \*\**P* < 0.01). (e) Heatmap showing coefficient of variation across all neutrophils for ten OMI and three morphological variables. Groups include unstimulated control, and neutrophils activated with *P. aeruginosa*, and *T. gondii* strains RH and M49. Data includes neutrophils from 3 distinct donors (Donor M, N and P, number of cells per donor in Table 1). Data in (b-e) collected 15 minutes after treatment.

Consistent with OMI results, a similar trend was observed in LC-MS-based time series redox ratio measurements (Fig 1j). Additionally, LC-MS analysis at later timepoints showed a progressive increase in the abundance of NADP+ at 30 minutes and 60 minutes after PMA treatment (Fig 1g, Table 3). In addition, analysis in the same subject showed NADPH levels decreased at 30 minutes and 60 minutes after PMA treatment, aligning with the optical redox ratio and NAD(P)H mean lifetime trends (Fig 1g).

We further investigated metabolic pathway-related metabolites at later time points - 30 minutes, and 60 minutes after PMA treatment. We observed a progressive rise in the PPP intermediates (Fig 1g). Additionally, glycolysis intermediates were found to be more abundant compared to unstimulated control conditions over the same time points (Fig 1g). Notably, we found a greater abundance of glutamate in activated neutrophils (Supp Fig 3a). Glutamine metabolism has been previously observed in human neutrophils^2,21^, and this observation is consistent with the possibility that activated neutrophils employ glutaminolysis as a supplementary mechanism for NADPH production, where the amino acid glutamine is oxidized to glutamate^22^. We also observed an increase in the abundance of methionine sulfoxide, particularly at 60 minutes post-PMA treatment (Supp Fig 3a, Table 3). This could indicate oxidation of methionine to methionine sulfoxide by ROS during oxidative burst, which is expected to increase around 60 minutes after activation. Similar oxidation of methionine residues have been previously reported in PMA-activated neutrophils^23^. Finally, PMA also resulted in an increase in the abundance of certain nucleotides including inosine, hypoxanthine, and IMP. (Supp Fig 3b).

To further characterize the OMI parameters in activated neutrophils, we employed a dimension reduction method, Uniform Manifold Approximate and Projection (UMAP) (Fig 2a). Along with the imaging parameters, morphological features were also included in the UMAP (Table 4), as neutrophils are known to change morphology upon activation. Except for a few donor-dependent variabilities, UMAP showed clear separation between control (unactivated) and PMA-activated neutrophils.

Lastly, to determine whether OMI can predict the activation status of neutrophils, we used machine learning to classify PMA-treated cells versus untreated cells at 15 minutes post-stimulation. A random forest classifier, developed based on NAD(P)H and FAD lifetime features along with morphological features (Table 4), achieved high sensitivity and specificity (AUC = 0.9) in predicting activation (Fig 2b). A random forest classifier constructed using NAD(P)H and FAD lifetime features exclusively also achieved high classification accuracy with AUC of 0.84 (Supp Fig 4d).

Overall, these results highlight how NAD(P)H lifetime and optical redox ratio can provide complementary insights into the use of metabolic pathways during neutrophil activation, here revealing an immediate shift toward an oxidative state upon activation in primary human neutrophils. Moreover, quantification of these metabolic changes in single cells by OMI at 15- minutes post-stimulation holds promise for predicting later neutrophil activation status.

### Optical metabolic imaging is sensitive to metabolic flexibility in activated neutrophils

To investigate the effect of inhibiting glycolysis and PPP in activated neutrophils, we first verified that PMA produces an oxidative burst by generating ROS through the NOX2 enzyme complex. For this, we inhibited NOX2 complex activity in PMA-stimulated neutrophils with diphenyleneiodonium (DPI)^24^ (Fig 3a). As expected, when the complex was inhibited, PMA did not produce an oxidative burst, confirming that ROS produced at NOX2 are the major contributors to oxidative burst in PMA-activated neutrophils (Fig 3b, Supp. Fig 5a). Consequently, DPI also inhibited NETosis (Supp Fig 1d).

We next sought to delve deeper into the metabolic pathways potentially responsible for the observed oxidized state of neutrophils within 15 minutes following activation. To do so, we selectively inhibited glycolytic and PPP pathways in activated neutrophils with the commonly used inhibitors 2-Deoxy-d-glucose (2-DG), a competitive inhibitor of glucose uptake, and 6- aminonicotinamide (6-AN), an NADPH-competitive inhibitor that blocks G6PD activity, respectively^3^ (Fig 3a).

First, the oxidative burst was assessed in the presence of metabolic inhibitions using a fluorescence indicator of ROS. Oxidative burst was diminished upon inhibition of both glycolysis and PPP pathways, underscoring the critical roles of these pathways in maintaining ROS production in activated neutrophils (Fig 3b, Supp Fig 5a). In contrast, inhibition of mitochondrial electron transport chain (ETC) complex III and complex IV with Antimycin A (AA) and sodium cyanide (NaCN) respectively^3^ did not hinder oxidative burst in PMA-activated neutrophils (Fig 3a, Supp Fig 5a). This result suggests an insignificant contribution of mitochondrial ROS to the oxidative burst under PMA stimulation. Notably, similar results were obtained when the neutrophils were pre-treated with the inhibitors before activation (Supp Fig 5a). Importantly, the treatments did not cause significant neutrophil death (Supp Fig 1b).

**Figure 5.**
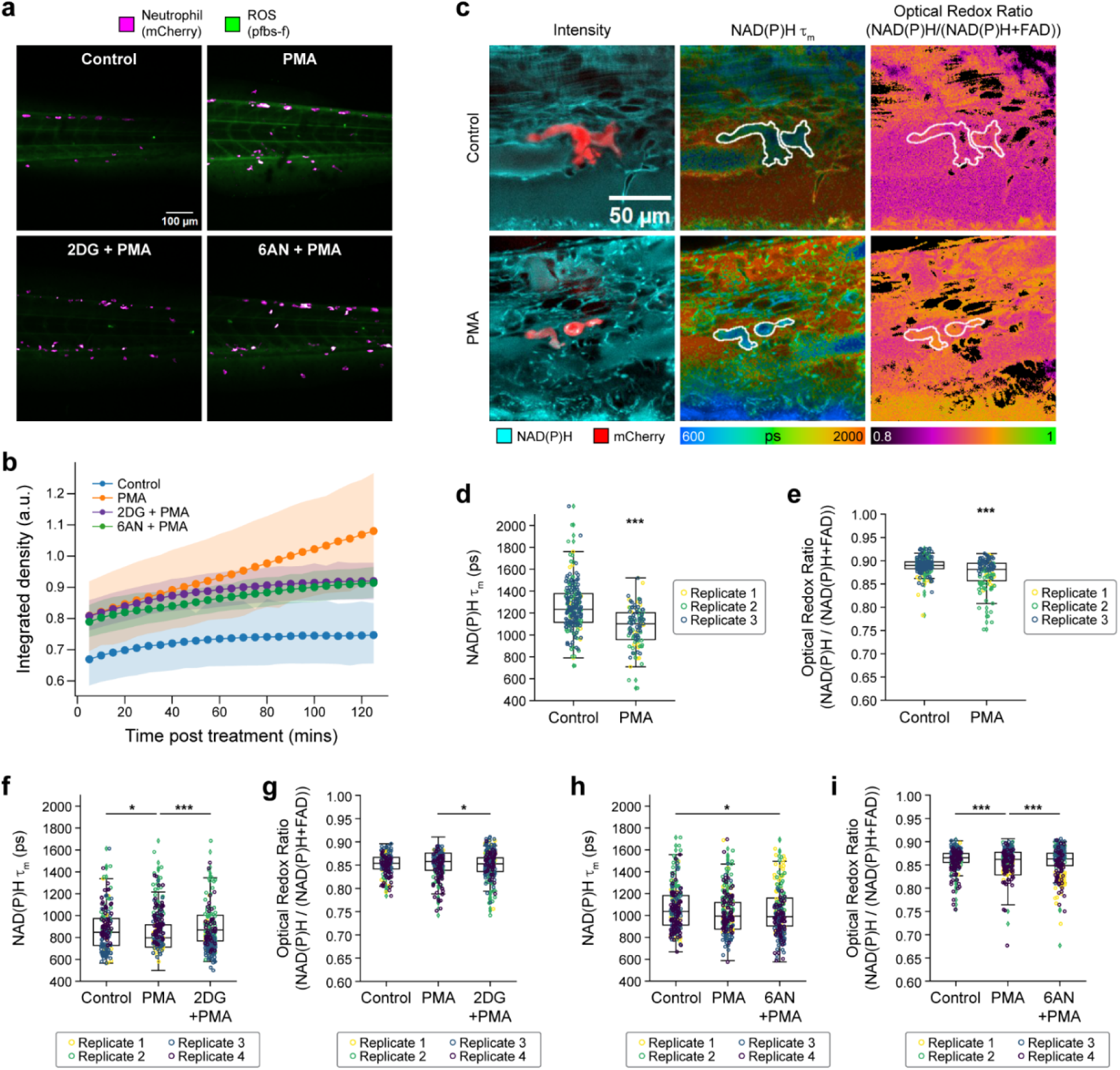
Metabolic rewiring of activated neutrophils *in vivo* in zebrafish larvae. (a) Representative fluorescence intensity of 1 mM pentafluorobenzenesulfonyl fluorescein (pfbs-f), (green) and mCherry (magenta) representing ROS (H_2_O_2_) and neutrophils, respectively, approximately 60 minutes post-stimulation in control, 50 nM PMA and PMA along with inhibitors 10 mM 2-DG and 1 mM 6-AN treated 3 day post-fertilization (dpf) zebrafish larvae and (b) corresponding quantification of pfbs-f fluorescence intensity integrated density indicating oxidative burst. (c) Left panel shows representative fluorescence intensity images of NAD(P)H fluorescence (cyan) and mCherry (red), NAD(P)H mean lifetime (middle) and optical redox ratio (right) of control (top) and PMA treated (bottom) 3 dpf zebrafish larvae. Single-cell quantification of (d) NAD(P)H mean lifetime and (e) optical redox ratio of control and PMA treated 3 dpf zebrafish larvae neutrophil from 3 independent replicates with 3–5 larvae per replicate per condition and 3–6 images per larvae (n = number of cells in Table 6). Single-cell quantification of (f) NAD(P)H mean lifetime and (g) optical redox ratio of control (vector = E3 media control), PMA and PMA with 2-DG treated 3 dpf zebrafish larvae neutrophil from 3 independent replicates with 3-5 larvae per replicate per condition and 3–6 images per larvae, (n = number of cells in Table 6). Single-cell quantification of (h) NAD(P)H mean lifetime and (i) optical redox ratio of control (vector = DMSO vector control), PMA and PMA with 6-AN- treated 3 dpf zebrafish larvae neutrophil from 3 independent replicates with 3–5 larvae per replicate per condition and 3–6 images per larvae, (n = number of cells in Table 6). For statistical tests in d-i, general linear models were fitted to data with an interaction term for replicate and treatment. Influence of replicate on the observed trend was insignificant except for e. All data besides (b) were collected approximately 60 minutes after treatment.

Second, we studied the metabolic response of activated neutrophils with targeted inhibition of glycolysis, PPP, and mitochondrial oxidative phosphorylation (Fig 3a, Fig 3c). Compared to the PMA-treated group, 2-DG caused an increase in NAD(P)H mean lifetime and decrease in optical redox ratio and decrease in LC-MS-based redox ratio, suggesting a decrease in NAD(P)H with upper glycolysis inhibition (Fig 3d-f, Supp Fig 5d-f). Although the NAD(P)H mean lifetime of activated neutrophils treated with 2DG was higher than the PMA-stimulated group alone, it was still lower than that of unstimulated controls. This reduction in NAD(P)H mean lifetime with 2- DG compared to control aligns with previous observations in activated human T cells *in vitro*^13^ and in macrophages *in vivo*^25^. This suggests that inhibition of glycolysis in activated neutrophils may lead to a transition towards an alternate route, such as glutaminolysis, to generate free NADPH. Glutaminolysis has been previously reported to fuel the TCA cycle by generating α- ketoglutarate in neutrophils in glucose-limited conditions to meet their energy demands^26–28^. Moreover, glutaminolysis and TCA generate free NADPH and NADH respectively^2,5,29,30^, which could contribute to the observed decrease in NAD(P)H mean lifetime. To verify this, we investigated the metabolites involved in these pathways (Fig 3f, Supp Fig 6a-b, Table 5). We found that compared to stimulation with PMA, treatment with 2-DG resulted in a significant reduction of metabolites associated with glycolysis and PPP alongside elevated levels of metabolites linked with glutaminolysis and the TCA cycle, consistent with neutrophil use of glutaminolysis and the TCA cycle under glycolytic inhibition.

Inhibition of PPP with 6-AN resulted in a greater decrease in NAD(P)H mean lifetime and optical redox ratio compared to the inhibition of glycolysis with 2-DG in three out of four donors (Fig 3d-e, Supp Fig 5d-e), highlighting donor-specific variability with 6-AN treatment. Additionally, inhibition of PPP alone might alter the glycolytic pathway. PMA-stimulated neutrophils with 6-AN treatment displayed higher levels of the glycolysis-associated metabolites compared to both the PMA-stimulated and 2-DG-treated conditions (Fig 3f). Notably, 2-DG treatment, but not 6-AN, resulted in a decrease in ATP levels in activated neutrophils, consistent with the use of glycolysis in energy production in activated neutrophils (Supp Fig 6b).

Inhibition of the NOX2-complex by DPI in PMA-stimulated neutrophils (PMA+DPI), decreases the NAD(P)H mean lifetime compared to control conditions (Fig. 3d, Supp Fig 5d). This suggests that in the PMA+DPI treatment, while DPI impedes oxidative burst by inhibiting the NOX2-complex, PMA still triggers the generation of free NAD(P)H, as observed above with PMA alone. However, since DPI inhibits NOX2, the generated NADPH is not consumed, thus leading to a further increase in optical redox ratio (Fig. 3e). Consequently, LC-MS analysis also revealed an increase in glycolytic and PPP intermediates in activated neutrophils irrespective of DPI treatment, compared to unstimulated control (Fig. 1g). These findings align with the greater abundance of NADPH and lower abundance of NADP+ and FAD in PMA+DPI conditions compared to the PMA-alone conditions at 30mins and 60 mins after treatment (Fig 1g). Additionally, DPI did not alter the ATP levels compared to PMA-alone conditions (Supp Fig 6b). These results highlight the sensitivity of OMI to neutrophil metabolic alterations even when neutrophil functions are impaired.

Although inhibition of mitochondrial complexes did not impede the oxidative burst, the mitochondrial complex IV inhibitor NaCN plus the activator PMA induced a significant decrease in NAD(P)H mean lifetime and increase in optical redox compared to PMA alone (Supp Fig 5b- e). These are consistent with anticipated effects of the ETC inhibitors, and indicate the presence of competent mitochondrial respiratory chain complexes.

OMI cannot distinguish between NADH and NADPH since their spectral properties overlap^31^ . To assess the contribution of NADH and NADPH to the optical imaging readouts, we performed paired (neutrophils from the same donor on the same day) OMI and LC-MS measurements of activated neutrophils and activation with metabolic inhibitors (Supp Fig 7a-b). We found that both the NAD(P)H fluorescence intensity and optical redox ratio better correlated with the NADPH-based redox ratio (Fig 3g-h).

Taken together, the results indicate that activated neutrophils predominantly depend on glycolysis and PPP for their functional requirements. However, they also exhibit metabolic flexibility in adapting to their environment and OMI is highly sensitive to these dynamic metabolic changes.

### Live pathogens induce rapid and heterogenous metabolic response in neutrophils

To investigate the response to a physiologically relevant stimuli, we cocultured primary human neutrophils with two distinct pathogens, *Pseudomonas aeruginosa* (*P. aeruginosa*) and *Toxoplasma gondii* (*T. gondii*), and then measured oxidative burst to confirm activation and metabolic imaging to identify metabolic rewiring.

*P. aeruginosa* is a gram-negative, opportunistic bacteria associated with serious clinical pathologies in immunosuppressed patients and in diseases like cystic fibrosis, pneumonia, and sepsis^32^. Along with macrophages and lymphocytes, neutrophils play an important role in the host defense against this pathogen^33^. Adaptations in neutrophil metabolism for bacterial killing are not completely understood^27^. *P. aeruginosa* induced an oxidative burst, when cocultured with neutrophils, that started to increase around 50 minutes after exposure, compared to unstimulated control (Fig 4a, Supp Fig 8a).

While studies indicate that neutrophils might play a role in mitigating infection caused by *T. gondii*, a protozoan parasite, the response mechanism is not well understood^34–36^. To investigate the neutrophil activation and any metabolic remodeling that may occur, we exposed primary human neutrophils to two strains of *T. gondii* – type I-RH and type II- M49. The two *T. gondii* strains induced an oxidative burst that started to increase around 30-40 minutes of exposure and leveled off after ∼60 minutes of exposure. The less virulent M49-*T. gondii* strain induced somewhat less oxidative burst at later time points compared to the RH- *T. gondii* strain, perhaps indicating diminished response (Fig. 4a, Supp Fig. 8a). Overall, PMA elicited a higher level of oxidative burst compared to *T. gondii*. A similar increase in neutrophil NETosis frequency by PMA compared to the RH-*T. gondii* strain was previously observed ^35^.

Next, we investigated whether infection with these pathogens caused metabolic remodeling in neutrophils within 15 minutes of exposure (Fig 4b-d and Supp Fig 8b-c). However, the extent and alteration in OMI features induced by the three pathogens varied across pathogen and donor. Neutrophils from donors M and P showed decreases in NAD(P)H mean lifetime and increases in optical redox ratio with *P. aeruginosa* (Fig 4b-c and Supp Fig 8b-c). Yet, neutrophils from donor N did not display any change in NAD(P)H mean lifetime with *P. aeruginosa* and M49-*T. gondii*, but RH-*T. gondii* resulted in decreased NAD(P)H mean lifetime (Fig. 4c and Supp Fig 8b). This indicates that OMI is sensitive to donor-dependent metabolic adaptation of neutrophils under a variety of physiological stimulations.

Taking advantage of single cell metabolic features, we performed a heterogeneity analysis (coefficient of variation) between neutrophils at 15 minutes of exposure (Fig. 4e). The result shows the least variations in OMI and morphological parameters for the unstimulated neutrophils compared to pathogen-stimulated neutrophils. Conversely, the less virulent M49-*T. gondii* exhibited the most heterogeneity between neutrophils across all donors (Fig 4e, Supp. Fig 9). These results indicate that live pathogens elicit a variable metabolic response between neutrophils and human donors as detected by OMI.

### Neutrophil metabolic rewiring *in vivo* recapitulates *in vitro* results

To verify whether the metabolic requirements in neutrophils activated *in vitro* might translate to a more complex system, next we investigated the *in vivo* neutrophil response in zebrafish (*Danio rerio*) larvae. The zebrafish model has been used previously to monitor neutrophil function^37,38^, and several functional, behavioral, and morphological characteristics are conserved between mammalian and zebrafish neutrophils^38,39^. Neutrophils labeled with mCherry were imaged in live, three days post-fertilization (dpf) zebrafish larvae located in the caudal hematopoietic tissue (CHT) due to the high concentration of neutrophils in that region^40^. Simultaneous imaging of the mCherry-labeled neutrophils along with NAD(P)H and FAD captured metabolic information of neutrophils *in vivo*, a technique previously employed to study macrophages in zebrafish larvae^25^.

Neutrophil activation was induced by bathing larvae in 50 nM PMA for 2 hours. The activation of neutrophils *in vivo* was verified by a burst of extracellular ROS using pentafluorobenzenesulfonyl fluorescein (pfbs-f), a fluorescent probe for hydrogen peroxide^41^, approximately 60 minutes after the addition of PMA to the bath (Fig 5a-b). Oxidative burst was diminished when PMA-activated zebrafish larvae were exposed to 2-DG and 6-AN to block glycolysis and PPP, respectively (Fig 5a-b), similarly as *in vitro*. Next, we performed OMI of neutrophils in zebrafish larvae under similar conditions (Fig 5c). We found that PMA activation caused a significant decrease in NAD(P)H mean lifetime (Fig 5d), which aligns with the results observed in primary human neutrophils *in vitro* (Fig 1d). The optical redox ratio also decreased with PMA, suggesting NADPH oxidation at NOX2 since the data was acquired over 90 minutes after activation (Fig 5e), consistent with later time points in primary human neutrophils *in vitro* (Fig 1e).

Additionally, we found that glycolysis inhibition with 2-DG (PMA + 2DG) caused a significant increase in NAD(P)H mean lifetime and decreased optical redox ratio compared to PMA alone (Fig 5f-g), again in agreement with the *in vitro* results (Fig 3d-e). Interestingly, PPP inhibition with 6AN (PMA + 6AN) caused no change in NAD(P)H mean lifetime and an increase in the optical redox ratio compared to PMA alone (Fig 5h-i), in contrast to the decrease in NAD(P)H mean lifetime and optical redox ratio in primary human neutrophils *in vitro* with PMA + 6ANcompared to PMA alone (Fig 3d-e). This discrepancy may be due to the different timing of these measurements *in vitro* (15 minutes post-stimulation) and *in vivo* (∼60 minutes post-stimulation).

Taken together, these *in vivo* studies suggest that neutrophil metabolic responses to stimulation were reproducible within both *in vitro* and *in vivo* settings, and that OMI can be used to quantify these metabolic responses. Additionally, while functional assays like oxidative burst measure extracellular ROS produced in the whole larvae, the 3D imaging capabilities of OMI provide sensitivity to metabolic changes specifically within the neutrophil population.

## Discussion

Neutrophils are crucial for defending the host against pathogens, and their dysregulation can lead to significant harm. In cases of severe bacterial or fungal infections, there is a need to improve overall neutrophil activity to control infection. Conversely, conditions such as chronic inflammatory diseases can exhibit excessive neutrophil activation, requiring measures to decrease neutrophil activity. Additionally, in situations like sepsis, cancer, or autoimmune diseases, neutrophils deviate from their normal function to assume a pathogenic role that sometimes contributes to tissue damage, necessitating restoration of their regular function. This intricate balance of neutrophil activity in various diseases makes them a promising target for therapeutic intervention.

As metabolism is linked with cellular function, there is a growing focus on neutrophil metabolism and its impact on host outcomes across different disease contexts^2^. In this work we demonstrate rapid metabolic remodeling in which neutrophils transition to a more reduced state within minutes of activation, prior to the onset of effector functions like oxidative burst and NETosis. This is demonstrated by the decrease in NAD(P)H mean lifetime and increase in optical redox ratio, which is concurrent with an increase in intermediates of glycolysis and PPP measured in our LC-MS results. These findings align with previous studies showing the importance of these pathways in neutrophil activation^3,4^. The fast metabolic shifts support the rapid cellular events that occur in neutrophils upon stimulation^42^. For example, PMA causes Protein kinase C activation and translocation to the cell membranes within two to five minutes and release of granules within a minute^19^.

Moreover, the non-destructive nature of OMI allowed real time measurement of metabolic changes in activated neutrophils at the single cell level, revealing how the initial reduced redox state, due to the increase in NAD(P)H production, gradually shifts towards an oxidized state with the onset of oxidative burst on a population level. Our LC-MS results show a decrease in NADH and NADPH and an increase in NADP^+^ over time after PMA treatment (Fig. 1g). This shift could be due to the formation of the NOX2-complex, where NADPH binds to NOX2 and loses electrons, transitioning into the non-fluorescent oxidized state NADP^+^ (Supp. Fig 4b). In addition, when electrons are transported through NOX2-complex, FAD is reduced to non-fluorescent FADH2 by gaining two electrons from NADPH and is subsequently oxidized back to FAD in a two-step process, further contributing to the oxidized state during the later NOX2- derived ROS generation. As equilibrium is reached in the process of electron transfer between NADPH and FAD, a steady redox state may be achieved. The duration over which the optical redox ratio of PMA-activated neutrophils returned to the levels of unstimulated control (before further decrease) varied among three separate donors, ranging from 15 to 30 minutes after activation, indicating variability between donors. These results demonstrate that, in addition to the immediate remodeling of metabolic pathways associated with activation, OMI is also sensitive to NADPH–NOX2 binding and subsequent electron transfer through the activated NOX2 enzyme systems. Moreover, the temporal trend observed indicates the lag between stimulation and NOX2 enzyme complex assembly and activation. A similar initial increase followed by a decrease in NAD(P)H intensity after the generation of oxidized species has been previously observed^43,44^.

While metabolic inhibitors support the use of PPP and glycolysis in activated neutrophils, our results also highlight the metabolic flexibility of neutrophils as recently suggested^4,6^. For example, our LC-MS metabolite analysis suggests inhibition of glycolysis and PPP could cause alterations in other pathways like glutaminolysis. Although not investigated in this study, previous work has shown that neutrophils use gluconeogenesis, glycogenesis, and fatty acid metabolism, especially in glucose-limited conditions^6,22,45^. Additionally, metabolic response to mitochondrial inhibition suggests the presence of functioning mitochondrial respiratory chain complexes that could contribute to neutrophil function. We found a decrease in NAD(P)H mean lifetime and increase in optical redox ratio with inhibition of mitochondrial complexes in activated neutrophils (Supp Fig 5b-e). This supports the evolving research on the importance of mitochondrial metabolism in neutrophils, especially in maintaining mitochondrial membrane potential (Δψ_m_) which is essential for chemotaxis, a processes by which neutrophils migrate to the inflammation site^2,5^. Moreover, a previous study found that in neutrophils, Complex III maintains Δψ_m_ by receiving electrons from glycolysis-derived glycerol-3-phosphate (G3P), which is subsequently oxidized to dihydroxyacetone phosphate (DHAP) in the process ^46^. The same study also showed that inhibition of Complex III by AA resulted in lower mitochondrial membrane potential in neutrophils. In fact, we found a higher abundance of G3P when PMA- stimulated neutrophils were treated with AA, which may be due to a decrease in its oxidation (Fig 3f). Thus, besides maintaining membrane potential, mitochondria can also support glycolytic flux. This observation provides further evidence for the ongoing investigation of mitochondrial metabolism in neutrophils^47^.

Our study extends its findings to pathogen-induced activation, demonstrating that live pathogens induce similar metabolic rewiring as synthetic stimuli. While the metabolic response of neutrophils to bacterial infections has been well characterized, their response to *T. gondii* is understudied. Previous studies have shown that *T. gondii* (RH strain) infection activates human primary neutrophils, leading to NETosis and cytokine production, and subsequent adaptive immune response^35^. Conversely, another study demonstrated the detrimental effect of *T. gondii* on the inflammatory response of neutrophils^36^. Lastly, neutrophils were shown to release interferon-γ to confer resistance against *T. gondii* infections in the central nervous system of mice^48^. Thus, while previous studies have demonstrated neutrophil effector function and cytokine production with *T. gondii* infection, our work demonstrates a rapid metabolic response to these parasites to better characterize neutrophil metabolic rewiring. Both *P. aeruginosa* and *T. gondii* induce an oxidative burst with OMI changes suggesting metabolic alterations in glycolysis and PPP in neutrophils. A decrease in NET formation has been previously observed when neutrophils were pretreated with 2DG and DPI, prior to infection with the RH-*T. gondii* strain. This further supports our results that indicate a metabolic switch in *T. gondii* activated neutrophils. Notably, the RH-*T. gondii* caused a greater change in optical redox ratio and NAD(P)H mean lifetime compared to both *P. aeruginosa* and the M49-*T. gondii* strain, suggesting a more pronounced response of neutrophils to that infection. However, the measurements fluctuate between donors, eliciting variable metabolic response to the bacterial versus protozoan infection. Finally, OMI captured this heterogeneous response to pathogen activation across single neutrophils. Specifically, we observed higher metabolic heterogeneity in neutrophils infected with the M49 *T. gondii* strain compared to the RH-*T. gondii* strain and *P. aeruginosa*. This could be because differences between the *T. gondii* strains extend beyond virulence to include distinct kinetics, dynamics, and replication rates^49^. The RH *T. gondii* strain causes host mitochondria to localize around parasite-containing vacuoles due to a mitochondria association factor (MAF), which the M49 *T. gondii* strain lacks^50^. MAF in the RH *T. gondii* strain may also correlate with increased mitochondrial respiration in infected cells and a more reduced optical redox ratio^51^. Therefore, the stronger overall metabolic response to the RH strain likely describes the decreased heterogeneity between neutrophils compared to the M49 strain.

To validate *in vitro* findings, neutrophil response in zebrafish larvae was investigated using OMI. The optically transparent nature of zebrafish along with the development of transgenic zebrafish with mCherry labelled neutrophils makes them ideal for measuring real-time metabolic changes using optical imaging. Consistent with *in vitro* results, PMA-induced activation leads to metabolic changes indicative of increased glycolysis and PPP. While PPP inhibition did not induce significant lifetime changes perhaps due to longer exposure, glycolysis inhibition *in vivo* mirrors the metabolic alterations observed *in vitro*, underscoring the reproducibility of neutrophil responses measured by OMI across model systems.

There are certain caveats to this study. First, alternative metabolic pathways like fatty acid oxidation, gluconeogenesis, glycogenesis, and the TCA cycle were not investigated in this work. Secondly, the inability of OMI to distinguish between NADH and NADPH limits the interpretation, although LC-MS and OMI show a greater correlation of the optical redox ratio to NADPH than NADH in activated neutrophils. Finally, *in vivo* measurements were performed on the neutrophils in the CHT region of the zebrafish larvae due to their limited mobility. However, their less mature characteristics might not completely capture the true metabolic responses of mature neutrophils. Also, the study would benefit from extension into physiologically relevant activation of neutrophils *in vivo*.

This is the first study to measure rapid single-cell metabolic changes in activated neutrophils both *in vitro* and *in vivo*. Additionally, machine learning-based models that were trained on single-cell OMI variables along with morphological features could classify activated neutrophils within 15 minutes of stimulation with high accuracy (AUC of ROC, 0.9). Therefore, OMI could be used as a rapid, label-free diagnostic, for example, to predict seriousness and mortality in COVID-19 as in to prior work based on cell surface markers of neutrophil activation in patient blood samples^52^. Along with providing insights into neutrophil biology that can help identify therapeutic targets, this method could provide a tool for non-destructive neutrophil health screening during neutrophil manufacturing for neutropenic patients^53^, neutrophil manufacturing from induced pluripotent stem cells^54,55^, or manufacturing of chimeric antigen receptor neutrophils for cancer immunotherapy^56^.

## Methods

### Isolation and culture of human peripheral blood neutrophils

For neutrophil isolation, peripheral blood was freshly collected from healthy human donors following protocols approved by the Institutional Review Board of the University of Wisconsin– Madison (2018–0103), and informed consent was obtained from all the donors. Peripheral blood was drawn into sterile syringes containing heparin. Neutrophils were isolated using the MACSxpress Whole Blood Neutrophil Isolation Kit (Miltenyi Biotec), following the manufacturer’s instructions within 1–2 hours of blood draw. Following isolation, erythrocyte depletion (Miltenyi Biotec) was performed according to the manufacturer’s instructions. Upon conclusion of the isolation, neutrophils were spun down according to the manufacturer’s instructions. Neutrophils were spun down at 300g for 5 minutes and resuspended in Roswell Park Memorial Institute (RPMI) 1640 Medium (Gibco) supplemented with 10% Adult Bovine Serum (ABS), and 1% penicillin-streptomycin and kept at 37°C under 5% CO2. Isolation purity was verified by flow cytometry using antibodies against neutrophil surface markers anti-human CD11b (Alexa Fluor 594) (Biolegend) and anti-human CD15 (SSEA-1)(Alexa Fluor 488) (Biolegend). Data were analyzed with FlowJo software. Isolated cells were >96% CD11b+ CD15+ (Supplementary Fig. 1a). All experiments were concluded within 6 hours of neutrophil isolation.

### Pseudomonas aeruginosa culture

*Pseudomonas aeruginosa* (PAK strain) expressing mCherry were cultured overnight then sub-cultured and grown until mid-exponential phase. For co-culture experiments, the bacteria were opsonized in pooled human serum (MP Biomedicals) for 30 minutes at 37°C then washed 3 times in PBS and resuspended in RPMI at 2x10^8^/mL.

### Toxoplasma gondii in vitro culture

*T. gondii* transgenic stains type II-ME49 and type I -RH tachyzoites expressing mCherry were used for *in vitro* experimental infections. Both strains were propagated in human foreskin fibroblast (HFF) cells cultured at 37°C under 5% CO2^57^. HFFs were cultured in Dulbecco’s modified Eagle’s medium (DMEM) (Gibco) with 10% Fetal Bovine Serum (FBS), 2 mM L-glutamine, and 1% penicillin-streptomycin (Sigma-Aldrich).

### Zebrafish husbandry

All protocols using zebrafish in this study have been approved by the University of Wisconsin– Madison Research Animals Resource Center (protocols M005405-A02). Adult zebrafish were maintained on a 14 hr:10 hr light/dark schedule. Upon fertilization, embryos were transferred into E3 media (5 mM NaCl, 0.17 mM KCl, 0.44 mM CaCl2, 0.33 mM MgSO4, 0.025 mM NaOH, 0.0003% Methylene Blue) and maintained at 28.5°C. Adult transgenic line Tg(mpx:mCherry) was used that labels neutrophils by driving the cytoplasmic expression of mCherry under the myeloid-specific peroxidase promoter^40^. The transgenic line was outcrossed to casper fish^58^ to generate transgenic line devoid of pigmentation to circumvent interference with multiphoton imaging.

### Ethics

Animal care and use were approved by the Institutional Animal Care and Use Committee of the University of Wisconsin and strictly followed guidelines set by the federal Health Research Extension Act and the Public Health Service Policy on the Humane Care and Use of Laboratory Animals, administered by the National Institute of Health Office of Laboratory Animal Welfare. All protocols using zebrafish in this study have been approved by the University of Wisconsin-Madison Research Animals Resource Center.

### Neutrophil activation and metabolic inhibition

For *in vitro* experiments, freshly isolated neutrophils cell suspension either in Eppendorf tubes or plated on P-selectin-coated plates were activated using 100nM phorbol myristate acetate (PMA) (Fisher Scientific), 5µg/L tumor necrosis factor-α (TNF-α) (Peprotech) and 20µg/L lipopolysaccharide (LPS) (Sigma Aldrich) for indicated time. Metabolic inhibitors were added to the cell suspensions in conjunction with PMA unless indicated otherwise at the following concentrations - diphenyleneiodonium (DPI) (Sigma Aldrich): 10µM; 6-aminonicotinamide (6- AN) (Sigma Aldrich): 5mM; 2-Deoxy-d-glucose (2-DG) (Sigma Aldrich): 100mM; Sodium cyanide (NaCN) (Fisher Scientific): 4mM; Antimycin A (AA) (Sigma Aldrich): 1µM.

For *in vivo* neutrophil stimulation, zebrafish larvae were exposed to 50 nM PMA (Fisher Scientific) up to 2 hours by immersion. For 2-DG treatment by immersion, 10 mM 2-DG (Sigma Aldrich) was added in conjunction with PMA. For 6-AN treatment by immersion, the zebrafish larvae were preincubated with 1 mM 6-AN (Cayman Chemicals) for 1 hour before adding PMA.

### Pathogen Infection

For OMI, 2x10^5^ freshly isolated human neutrophils were co-incubated with 2x 10^6^ opsonized *P. aeruginosa* to attain a co-culture ratio of 1:10 and plated about 15 minutes before imaging. For oxidative burst assay, neutrophils were plated 15 minutes before imaging and the bacteria were added right before imaging.

Similarly, 2x10^5^ freshly isolated neutrophils were infected with RH or ME49 *T. gondii* tachyzoites at multiplicity of infection of 2; and plated about 15 minutes before metabolic imaging. For oxidative burst assay, neutrophils were plated 15 minutes before imaging and the parasites were added right before imaging.

### *In vitro* oxidative burst assay

Oxidative burst was measured by imaging fluorescent ROS probe dihydrorhodamine 123 (DHR) (Invitrogen). Neutrophils were incubated for 15 minutes with 10µM DHR prior to plating. Activators and inhibitors were added right before imaging. ROS signal was measured on a Nikon Ti-2E widefield fluorescence microscope coupled to a SOLA light engine (380–660 nm, Lumencor). DHR was excited with 60 ms exposure time about 12% power (>20mW at sample) and through an excitation filter of 465–495nm and a 20x objective (Nikon Plan Apo λ, 0.75 NA). Emission was collected using a standard fluorescein isothiocyanate (FITC, 515-555 nm) filter. 4– 5 fields of view were collected per well every 5–30 minutes for 4 to 5 hours. All channels were imaged sequentially.

For oxidative burst assay with pathogen infection, additional mCherry was imaged with 40 ms exposure time about 15% power (>20mW at sample) using an excitation filter of 550–590nm and emission filter of 608-683nm. Samples were excited through a 40x objective (Nikon Plan Apo λ, 0.95 NA). All channels were imaged sequentially. 4–5 fields of view were collected per well every 5 minutes for 3 hours.

Intracellular DHR intensity was quantified by identifying cell regions using the phase contrast microscopy segmentation toolbox (PHANTAST)^59^ on FIJI^60^. First, the masks were generated from the brightfield images of each field of view per timepoint using PHANTAST (sigma=0.7; epsilon=0.05). Next, the masks were applied to the corresponding DHR images, to extract the intracellular DHR intensities.

### *In vivo* oxidative burst assay

For live *in vivo* imaging, the 3 dpf larvae were loaded in a zWEDGI device^61^ in E3 media without methylene blue and supplemented with 0.16 mg/mL Tricaine (tricaine methanesulfonate; Fisher Scientific) to anesthetize larvae. The head region was secured by adding 2% low gelling agarose (A9045, Sigma) to prevent drifting during imaging. Neutrophil activation with PMA and metabolic inhibition with 2-DG and 6-AN were performed as described above. To stain hydrogen peroxide, a 30 min pre-incubation with 1 µM pentafluorobenzenesulfonyl fluorescein (Santa Cruz, CAS 728912-45-6) by immersion was included. Time-lapse z-stack images were acquired at 5 µm steps every 5 minutes for 2 hours on a spinning disk confocal microscope, with a confocal scanhead (CSU-X; Yokogawa) on a Zeiss Observer Z.1 inverted microscope and an EMCCD Evolve-512 camera (Photometrics), with a Plan-Apochromat 10X/NA 0.3 air objective (Zeiss) and Zen software (Zeiss). Fluorescence intensity quantification was performed on FIJI^60^. Integrated density, which is the sum of pixel intensities from all the slices, was calculated per timepoint per field of view. Background pixels, i.e, pixels that were not part of the larvae were excluded from this calculation.

### NETosis assay

NET release was assessed by monitoring the accumulation of extracellular DNA over time, as described here^6^. In brief, neutrophils were seeded at a density of 4 × 10^4^ cells per well in a 96- well tissue culture plate coated with Cell-Tak (Corning) and centrifuged at 200g for 1 minute with minimal acceleration/deceleration. Next, to label extracellular DNA, Cytotox Green Reagent (Incucyte) was added to the culture medium at a dilution of 1:4,000, and images were captured at intervals of 10–30 minutes after stimulation using an Incucyte live cell imager under standard culture conditions (37°C, 5% CO_2_). NETs are indicated by the fluorescent signal outside of cells and were quantified through image analysis employing IncuCyte S3 Basic Analysis software.

### Metabolomics

For extracting intracellular metabolites, neutrophils were washed with PBS after removing the culture medium. Pelleted neutrophils were extracted using 150 μL of cold acetonitrile/methanol/water (40:40:20 v:v:v) of liquid chromatography–mass spectrometry (LC– MS) grade (for each 2 million cells), followed by centrifugation at 20,627g for 5 minutes at 4°C to eliminate any insoluble residue. Samples were dried under N_2_ flow the resuspended in LC– MS-grade water as loading solvent. The soluble metabolites were analyzed using a Thermo Q-Exactive mass spectrometer connected to a Vanquish Horizon Ultra-High Performance Liquid Chromatograph. Metabolites were separated on a 2.1 × 100mm, 1.7 μM Acquity UPLC BEH C18 Column (Waters) employing a gradient of solvent A (97:3 H2O/methanol, 10 mM TBA, 9 mM acetate, pH 8.2) and solvent B (100% methanol). The gradient used was: 0 min, 5% B; 2.5 min, 5% B; 17 min, 95% B; 21 min, 95% B; 21.5 min, 5% B. The flow rate was maintained at 0.2 ml min^−1^. Data acquisition was performed using full scan mode. Metabolite identification relied on exact m/z and retention time, which were determined using chemical standards. Data acquisition was conducted using Xcalibur 4.0 software and analysis on Maven. Data was processed by background subtraction, followed by fold change calculations. Replicates were averaged, expressed as relative to control condition, then log base 2 transformed (Log_2_ (Sample Abundance/Control Abundance)). Control is either unstimulated control condition or PMA stimulated condition as indicated. Metabolite heatmaps were prepared on GraphPad Prism version 10.0.0. Metabolic pathway analysis was performed by HumanCyc pathway online software^62^. For computing the LC-MS based redox ratios, the ion count values were converted to molar concentration using external calibration curves.

### Cytotoxicity assay

A cytotoxicity assay was performed using CellTox™ Green Kit (Promega, Catalog: G8743) following the manufacturer’s instructions for the endpoint measurement 2X Reagent Addition. Briefly, cells were plated at 10,000 cells / well determined using the described CellTox Green linear range assay. Treatments were made at 2X concentration and cells were treated as follows: PMA [100nM], TNFa [5µg/L], LPS[20µg/L], PMA [100nM] along with inhibitors- 6AN [5mM], 2DG [100mM], DPI [10µM], AA [1µM], NaCN [4mM]. Alongside untreated, DMSO and the provided Assay Toxicity Control were used for control conditions. Finally, the 2x CellTox Green Reagent was made (20µL of Cell Tox Green Dye + 10mL of Buffer) and added to the cells immediately following the 15min treatments. Plate was allowed to sit in the 37C 5% CO_2_ incubator for 15min prior to plate reading. Fluorescence was measured on a microplate reader (Infinite M1000, Tecan) with the following settings: excitation wavelength: 485nm, emission wavelength: 535nm, gain was set based on one of the Toxicity control wells, 10 reads were taken per well with 40µs integration time. Average and standard deviations for all wells within a condition are reported.

### Sample preparation for multiphoton imaging

For *in vitro* imaging, neutrophils were plated at 200,000 cells per 50 µl of medium on either 35mm glass-bottom dishes (MatTek) or 24-well glass bottom plates with #1.5 cover glass (Cellvis) that were coated with 5µg/ml P-selectin (Bio-Techne) by incubating overnight at 4°C. Neutrophils were plated about 15 minutes before imaging. During imaging, the samples were housed in a stage top incubator (Tokai Hit) to keep them at 37°C and under 5% CO_2_. The activators and inhibitors were either added to the cells prior to plating or after plating.

For live *in vivo* imaging, the 3 dpf larvae were loaded in a zWEDGI device^61^, as described above. Neutrophil activation with PMA and metabolic inhibition with 2-DG and 6-AN were performed as described above in E3 media without methylene blue and supplemented with 0.16 mg/mL Tricaine (tricaine methanesulfonate; Fisher Scientific) to anesthetize larvae. The head region was secured by adding 2% low gelling agarose (A9045, Sigma) to prevent drifting during imaging.

### Optical metabolic imaging

Two-photon FLIM was performed on a custom-made Ultima Multiphoton Imaging System (Bruker) that consists of an inverted microscope (TI-E, Nikon). The system is coupled to an ultrafast tunable laser source (Insight DS+, Spectra Physics Inc). The fluorescence lifetime data was acquired using time-correlated single-photon counting electronics (SPC -150, Becker & Hickl GmbH) and imaging was performed using Prairie View Software (Bruker). For *in vitro* imaging, NAD(P)H and FAD were sequentially excited using excitation wavelengths of 750 nm and 890 nm respectively, and the laser power at the sample was <10 mW. The samples were illuminated using a 40x objective lens (W.I./1.15 NA /Nikon PlanApo) with a pixel dwell time of 4.8 µs and frame integration of 60 s at 256×256 pixels and 2x zoom of maximum field of view (∼0.9 mm^2^). The photon count rates were maintained at 1–5 × 10^5^ photons/second to ensure adequate photon detection for accurate lifetime decay fits. A dichroic mirror (720 nm) and bandpass filters separated fluorescence signals from the excitation laser. Emission filters used were bandpass 460/80 nm for NAD(P)H and 500/100 nm for FAD. For co-culture experiments with mCherry labeled pathogens, mCherry was imaged sequentially with two-photon excitation of 1040 nm and emission was collected using bandpass 590/45nm. Fluorescence signals were collected on GaAsP photomultiplier tubes (H7422P-40, Hamamatsu, Japan). The instrument response function (IRF) was collected each day by recording the second harmonic generation signal of urea crystals (Sigma-Aldrich) excited at 890nm.

For *in vivo* imaging NAD(P)H, FAD and mCherry were imaged simultaneously by wavelength mixing as described previously^25,63^. Briefly, wavelength mixing was achieved by tuning the multiphoton laser to 750 nm (λ1), that excited NAD(P)H, and was delayed and collimated with a second fixed laser line of 1040 nm (λ2), that excited mCherry, to obtain spatial and temporal overlap of the two pulse trains at each raster-scanned focal point. This created a third 2-color, two-photon excitation given by λ3 = 2/ (1/λ1 + 1/λ2), used to excite FAD. Fluorescence signals from NAD(P)H, FAD and mCherry were collected using bandpass filters 466/40nm, 540/24 nm, and 650/45 nm respectively.

### Image analysis and single-cell quantification

Fluorescence lifetime data were analyzed on SPCImage software (Becker & Hickl). Photon counts at each pixel were enhanced by a 3x3 binning comprising of 9 surrounding pixels. To eliminate pixels with a low fluorescence signal, an intensity threshold was used. NAD(P)H has an open conformation in its bound state while it adopts a closed conformation in its free state and hence has a comparatively shorter fluorescence lifetime. Conversely, FAD has a long lifetime in the free state and a shorter lifetime in bound state. Thus, using an iterative parameter optimization to obtain the lowest sum of the squared differences between model and data (Weighted Least Squares algorithm), the pixel-wise NAD(P)H and FAD decay curves were fit to a biexponential model [I(t) = α_1_×exp(−t/τ_1_) + α_2_×exp(−t/τ_2_) + C)] convolved with the system IRF. Here, I(t) represent the fluorescence intensity measured at time t, α_1_, α_2_ are the fractional contributions and τ_1_, τ_2_ denote the short and long lifetime components, respectively. C accounts for background light. The goodness of the fit was checked using a reduced chi-squared value<1.0. The mean lifetime is the weighted average of the free and bound lifetimes (τ_m_ = α_1_×τ_1_ + α_1_×τ_2_) and is calculated for each pixel. To generate the intensity of NAD(P)H (I_NAD(P)H_) and FAD (I_FAD_), the area under the fluorescence decay curve at each pixel was integrated. The optical redox ratio at each pixel is given by [I_NAD(P)H_ / (I_NAD(P)H_ + I_FAD_)].

Automated whole cell masks were created from NAD(P)H intensity images using Cellpose 2.0 (model = cyto 2, radius = 30)^64^. Following this, the generated masks were manually edited on Napari^65^. Masks were excluded if they were (1) image boundary cells with <50% of the cell area within the image, (2) other immune cells evidenced by longer NAD(P)H mean lifetime, single lobed nuclei and smaller size, (3) blurry/moving cells, (4) pathogens identified from mCherry signal and (5) red blood cells evidenced by very short lifetime and small size. For *in vivo* images, single cell masks were generated using mCherry intensity and lifetime data was used to exclude masks as needed. All the processing and analysis of single-cell data was done using a custom python library Cell Analysis Tools that extracted OMI and morphological features for each cell mask^66^.

### Statistical analysis

Analysis of the OMI data was performed in Python, and graphs for all the figures were plotted in Python using the Seaborn data visualization library. Statistical analysis was performed in Jamovi^67^, with either Student’s t test or one-way analysis of variance (ANOVA) with a or Tukey’s post hoc test for multiple comparisons, chosen based on data features. P values have been indicated as follows: *p< 0.05; **p < 0.01; ***p < 0.001, with p < 0.05 chosen as the threshold for statistical significance. For *in vitro* OMI data, every data point represents a neutrophil. Statistical significance was determined using one-way ANOVA with post hoc Tukey’s test. (***P < 0.001; *P < 0.05). Error bars are 95% confidence interval unless indicated otherwise. For *in vivo* zebra larvae, biological repeats are defined as separate clutches of embryos collected on separate days. For statistical analysis, general linear models were fit to data in Jamovi^67^ using the GAMLj module. Every data point represents a neutrophil. An interaction term was added to the model where more than one experimental factor was present (replicate, treatment).

### Classification

Uniform Manifold Approximate and Projection (UMAP) and coefficient of variation z-score heatmaps were used to visualize clustering (Seaborn, Python). Random forest classifier was used for machine learning based classification (R). All ROC curves and confusion matrices were constructed from the test data sets using the model generated from the training data sets. The classifiers in Fig 2b and Supp. Fig 4d were trained and tested on data from donors 1-8, with a split ratio of train: 70%, test: 30% of cells.

## Author contributions

Conceptualization: RD*, MCS* Methodology: RD*, VM Investigation: RD*, VM, EB, TQ Formal analysis: RD*, GMG, AG, AK, JV Resources: GMG, MAG Funding acquisition: JF, AH, MCS* Writing – Original Draft Preparation: RD*, MCS* Writing – Review & Editing: RD*, VM, GMG, JF, AH, MCS*

* corresponding authors

## Acknowledgements

The authors would like to thank Dr. Laura Knoll for providing us with the *T. gondii* strains; Alicia Williams for editing the manuscript; and Jens C. Eickhofffor reviewing the statistical analysis; Matthew Stefely for visualizations and figure editing. We would also like to acknowledge Kelsey Tweed, Stephen Halada, Rebecca Schmitz, and Peter Rehani for their help with this work.

## Data availability

The LC-MS data generated in this study is being deposited in the metabolomics workbench database DRYAD. The m/z cloud database will be available, and all the dataset files can be downloaded upon publication.

---

**Supplementary Figure 1:**
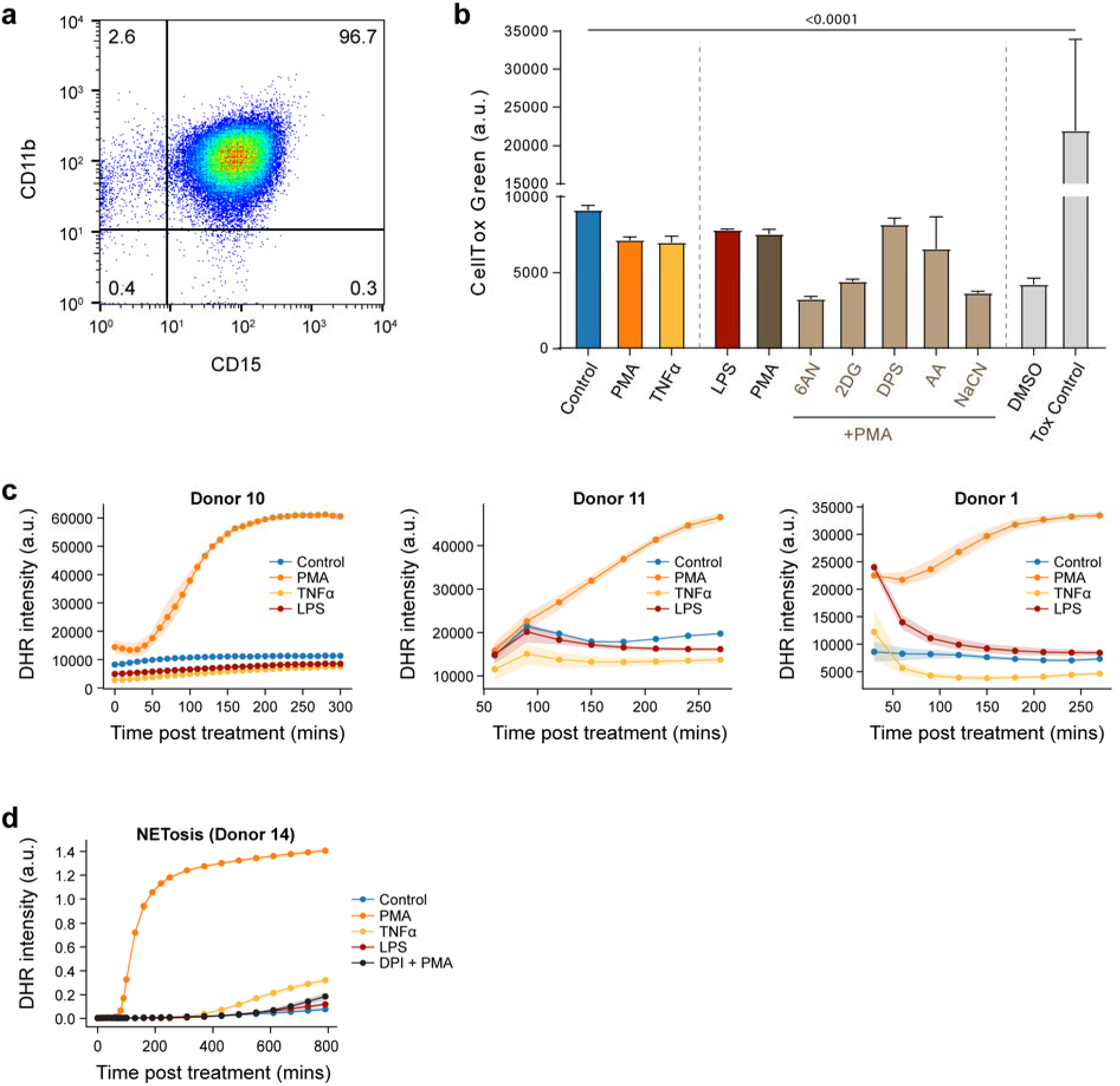
(a) Quantification of neutrophil isolation purity by flow cytometry using labels for CD11b and CD15. (b) Cytotoxicity assay of neutrophils using membrane-impermeable dye Celltox green. The conditions include unstimulated control, (100nM), LPS (20µg/L) and TNFα (5µg/L) treatment for 60 mins and PMA, PMA along with inhibitors 100mM 2DG, 5mM 6AN, 10µM DPI, and 1µM AA treatment for 15 mins. (c) Quantification of fluorescence intensity of DHR indicating intracellular ROS in unstimulated control and PMA (100nM), TNFα (5µg/L) and LPS (20µg/L) treated neutrophils. These are repeats of from 3 distinct donors (Donor 10, 11 and 1) compared to data presented in Fig 1a. (d) Quantification of NET release assay. A subset of this data was presented in Fig 1b.

**Supplementary Figure 2.**
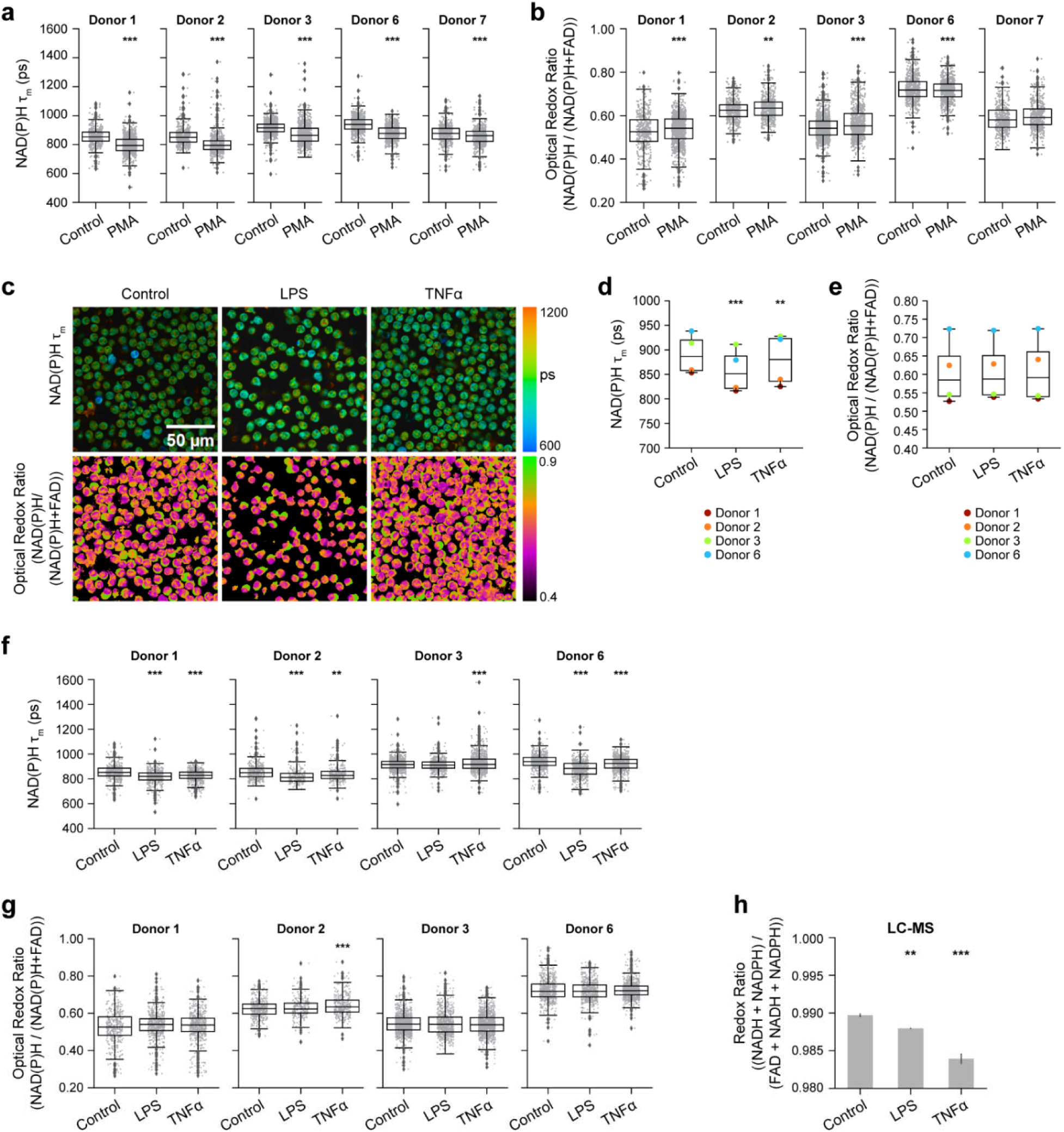
Single cell quantification of (a) NAD(P)H mean lifetime and (b) optical redox ratio of control and PMA (100nM) treated neutrophils separately plotted for 5 distinct donors (Donors 1, 2, 3, 6, 7) Single cell quantification for Donors 4-5 is presented in Supp. Fig 5d-e and for Donor 8 in Supp. Fig 7a. At least 5 images were acquired per condition and each data point is a single cell: number of cells in Table 1. Data presented in Fig 1d and 1e are derived from (a) and (b) respectively. (c) Representative images of NAD(P)H mean lifetime and optical redox ratio of control, LPS (20µg/L for 60 mins) and TNFα (5µg/L for 60 mins) treated neutrophils. Donor-level average of (d) NAD(P)H mean lifetime and (e) optical redox ratio of control, LPS and TNFα treated neutrophils from 4 distinct donors (Donors 1, 2, 3, and 6). Each point represents the average value for a single donor (n=number of cells across all donors, Table 1). Corresponding single cell quantification of (f) NAD(P)H mean lifetime and (g) optical redox ratio of control and LPS and TNFα treated neutrophils plotted separately for the 4 distinct donors where each point is a single cell. Data presented in (d) and (e) are derived from (f) and (g) respectively. (h) Redox ratio computed from molar concentration measured by LC-MS from 2 technical replicates for indicated conditions. The control from this data was presented in Fig 1f. The control data for Donor 1-3, 6 in (d-g) are also presented in (a-b) and Fig 1d-e. Statistical significance of differences between multiple conditions for data presented in (a-g) were tested using ANOVA with posthoc Tukey’s test (*** p< 0.001; ** p< 0.01; * p< 0.05). Error bars represent the 95% confidence interval.

**Supplementary Figure 3.**
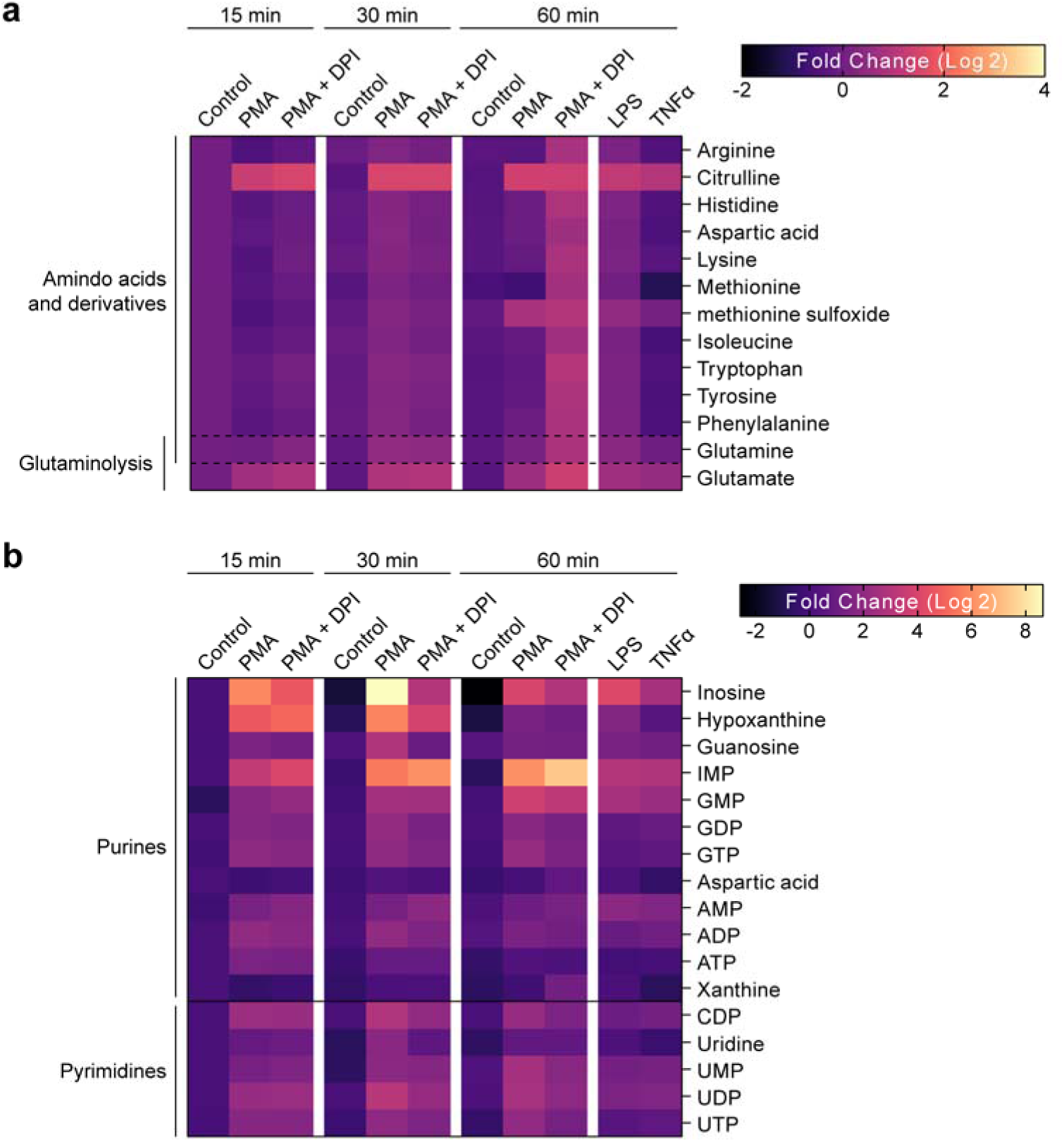
: Heat map representation of metabolomic variations across the unstimulated control, PMA treatment (100nM) and PMA together with NOX2 inhibitor DPI (10µM) treatment (PMS+DPI). Heatmap also includes stimulation with LPS (20µg/L) and TNFα (5µg/L). To align with the OMI conditions, metabolites were extracted at 15, 30 and 60 mins for all the conditions except for LPS and TNFα, which were extracted at 60 mins. Each metabolite abundance is normalized to the control abundance at 15 mins then log base 2 transformed (Log2 (measured Abundance/Control Abundance)) for each experimental condition. Metabolites have been grouped based on (a) amino acids and derivatives and glutaminolysis, and (b) nucleotides. The results of the significance tests are presented in Table 3.

**Supplementary Figure 4.**
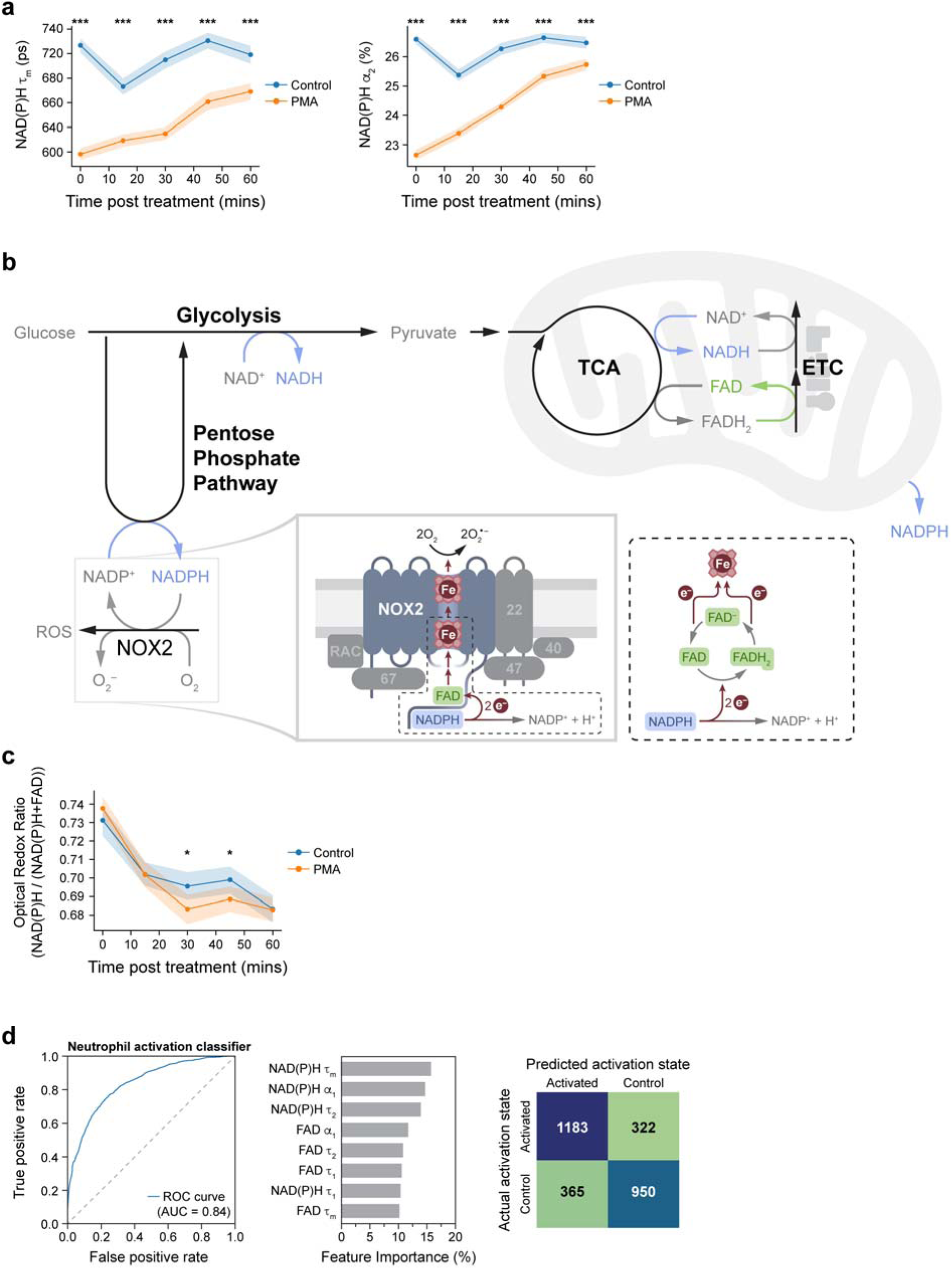
: Single cell quantification of (a) NAD(P)H mean lifetime (left), and bound NAD(P)H α_2_ percentage (right) of control and PMA (100nM) treated neutrophils acquired from at least 5 images per condition every 15 mins from 0 mins (after addition of PMA) to 60 mins post-treatment. Each point represents the average for all cells at the indicated timepoint, number of cells in Table 2 and error bars represent the 95% confidence interval. This is the second repeat on neutrophils from a distinct donor (Donor 9) compared to data presented in Fig 1g. (b) Schematic of major metabolic pathways. Inset shows the route of electron transfer in the activated NOX2 enzyme complex consisting of integral transmembrane proteins NOX2, p22phox (22) and associated cytosolic proteins p40phox (40), p47phox (47), p67phox (67) and RAC (RAC1 or RAC 2). NADPH in the cytosol donates 2 electrons which are transferred by FAD to two heme (Fe) groups and ultimately to molecular oxygen to generate superoxide (2O_2_ ^-^). In this process, NADPH is reduced to NADP+ while FAD cycles between FADH2 (by receiving 2 electrons) and FAD (by donating 2 electrons in a 2-step process). **(c)** Optical redox ratio of control and PMA treated neutrophils corresponding to (a). Each point represents the average for all cells at the indicated time point and error bars represent the 95% confidence interval. This is the second repeat on neutrophils from a distinct donor (Donor 9) compared to data presented in Fig 1h. (d) ROC curve and AUC of random forest model to classify neutrophil activation state using only OMI features (subset of features presented in Fig 2b) and corresponding feature importance and confusion matrix. The classifier was trained and tested on data from donors 1-8, with a split ratio of train: 70%, test: 30% of cells (number of cells in Table 1)

**Supplementary Figure 5.**
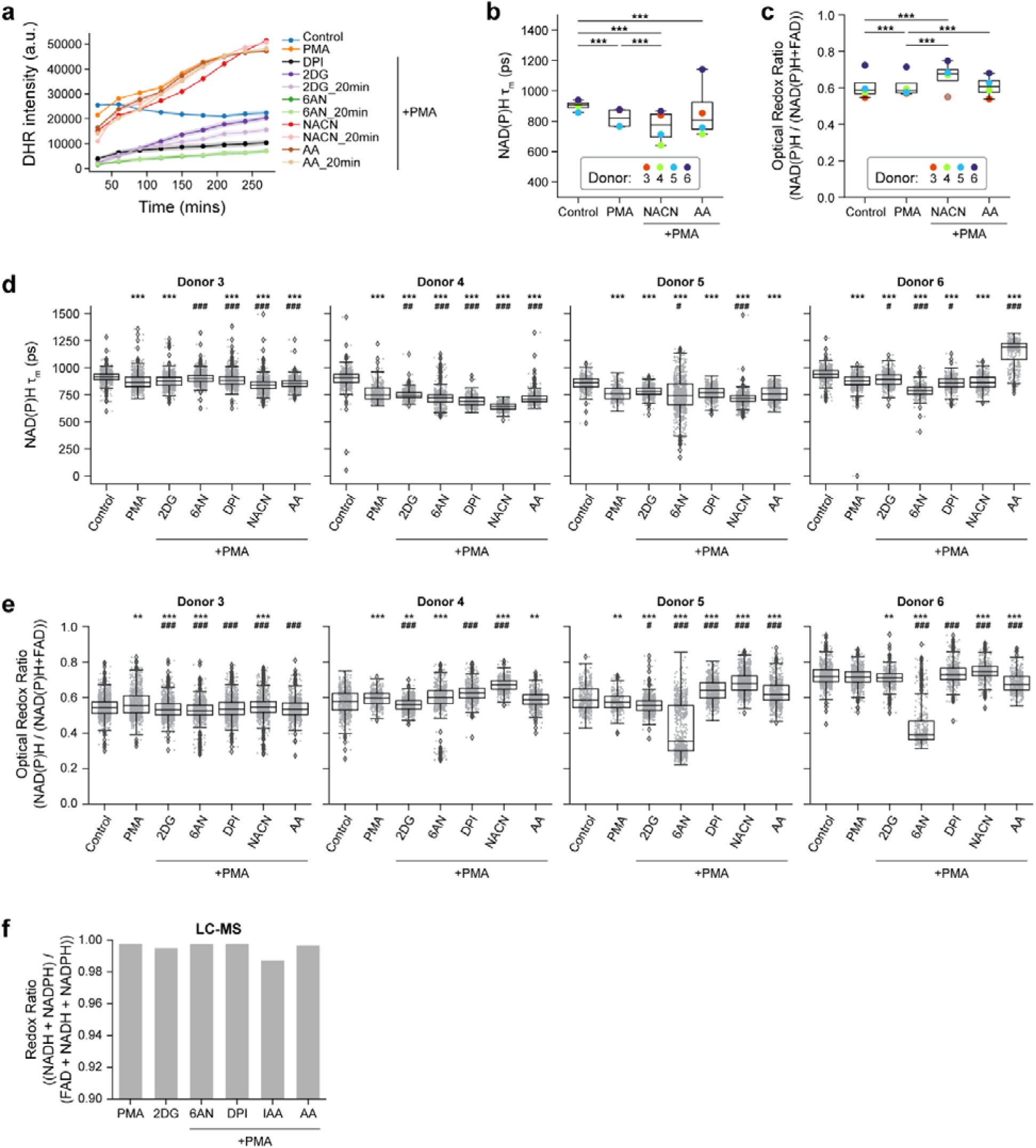
: (a) Quantification of fluorescence intensity of Dihydrorhodamine 123 (DHR) indicating intracellular ROS in unstimulated control, PMA treated, PMA along with 100mM 2DG, 5mM 6AN, 10µM DPI, and1µM AA treated neutrophils for 15 minutes. Inhibitor labels ending with ‘20’ indicate pre-incubation with the inhibitor for 20 minutes before PMA treatment. A subset of conditions are repeats of Fig 3b but on neutrophils from a different donor (Donor 12). (b) NAD(P)H mean lifetime and (c) optical redox ratio of control, PMA treated, and PMA plus inhibitor (NaCN, AA) treated (15 mins) neutrophils from 4 distinct donors (Donor 3- 6). Each dot represents the average across all cells per donor (n=number of cells across all donors, Table 1). Corresponding single cell quantification of (d) NAD(P)H mean lifetime and (e) optical redox ratio separately plotted for the 4 distinct donors (Donor 3-6) where each point is a single cell. Data presented in (b) and (c) are derived from (d) and (e) respectively.Symbols for statistical significance in d-e: * indicates comparison to control, # indicates comparison to PMA. (f) Redox ratio computed from LC-MS measurements. Statistical significance of differences between multiple conditions for data presented in (b-e) were tested using ANOVA with posthoc Tukey’s test (***/### p< 0.001; **/## p< 0.01; */# p< 0.05). Error bars represent the 95% confidence interval

**Supplementary Figure 6.**
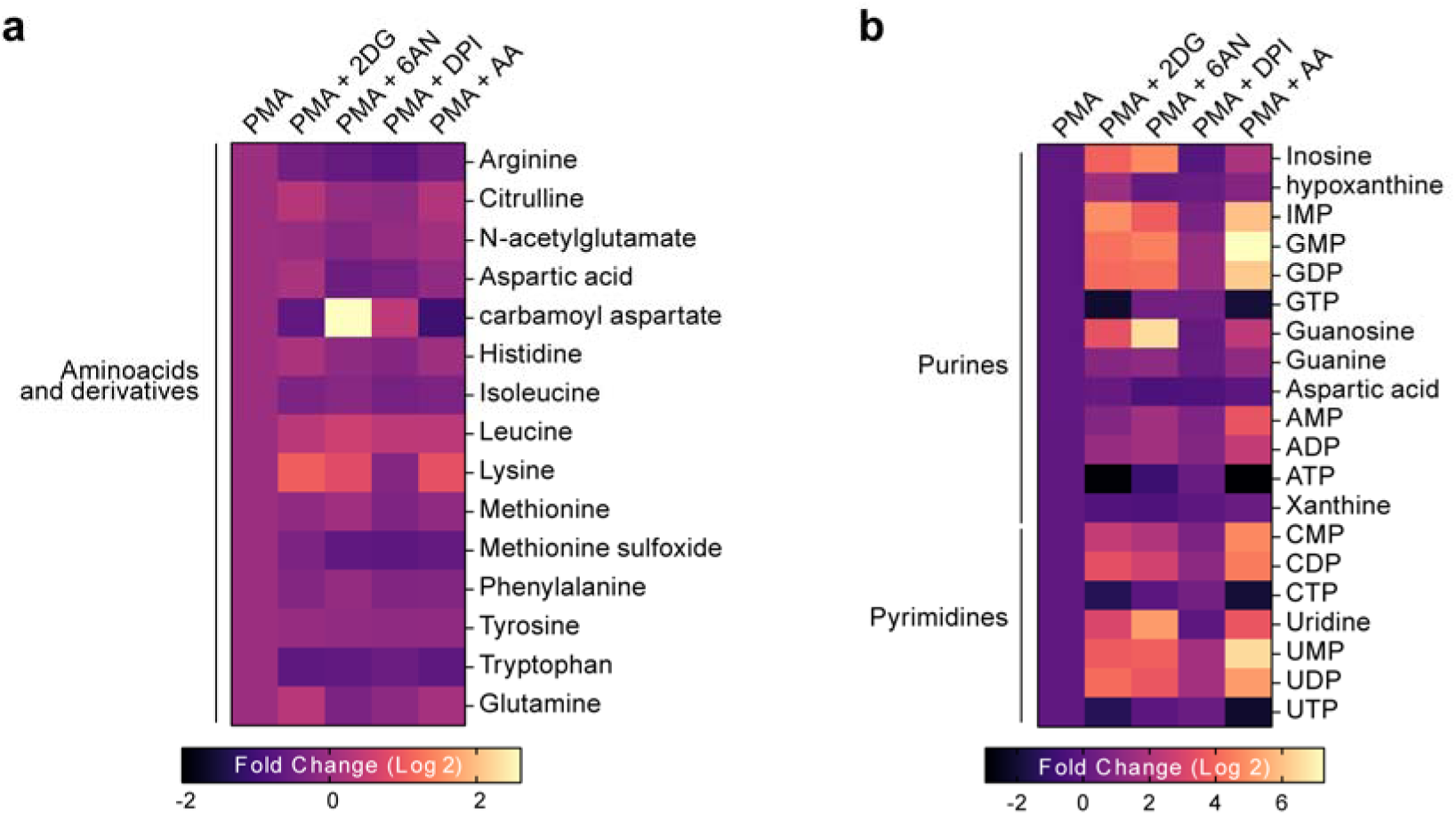
: Heat map representation of metabolomic variations across PMA (100nM) treatment and PMA together with 100mM 2DG (PMS+2DG), 5mM 6AN (PMA+6AN), 10µM DPI (PMA+DPI), and 1µM AA (PMA+AA) (Donor 13). To align with the OMI conditions, metabolites were extracted at 15 minutes. Each metabolite abundance is normalized to the control abundance then log base 2 transformed (Log_2_ (measured Abundance/Control Abundance)) for each experimental condition. Metabolites have been grouped based on (a) amino acids and derivatives, and (b) nucleotides (c). Result of significance test is presented in Table 5

**Supplementary Figure 7:**
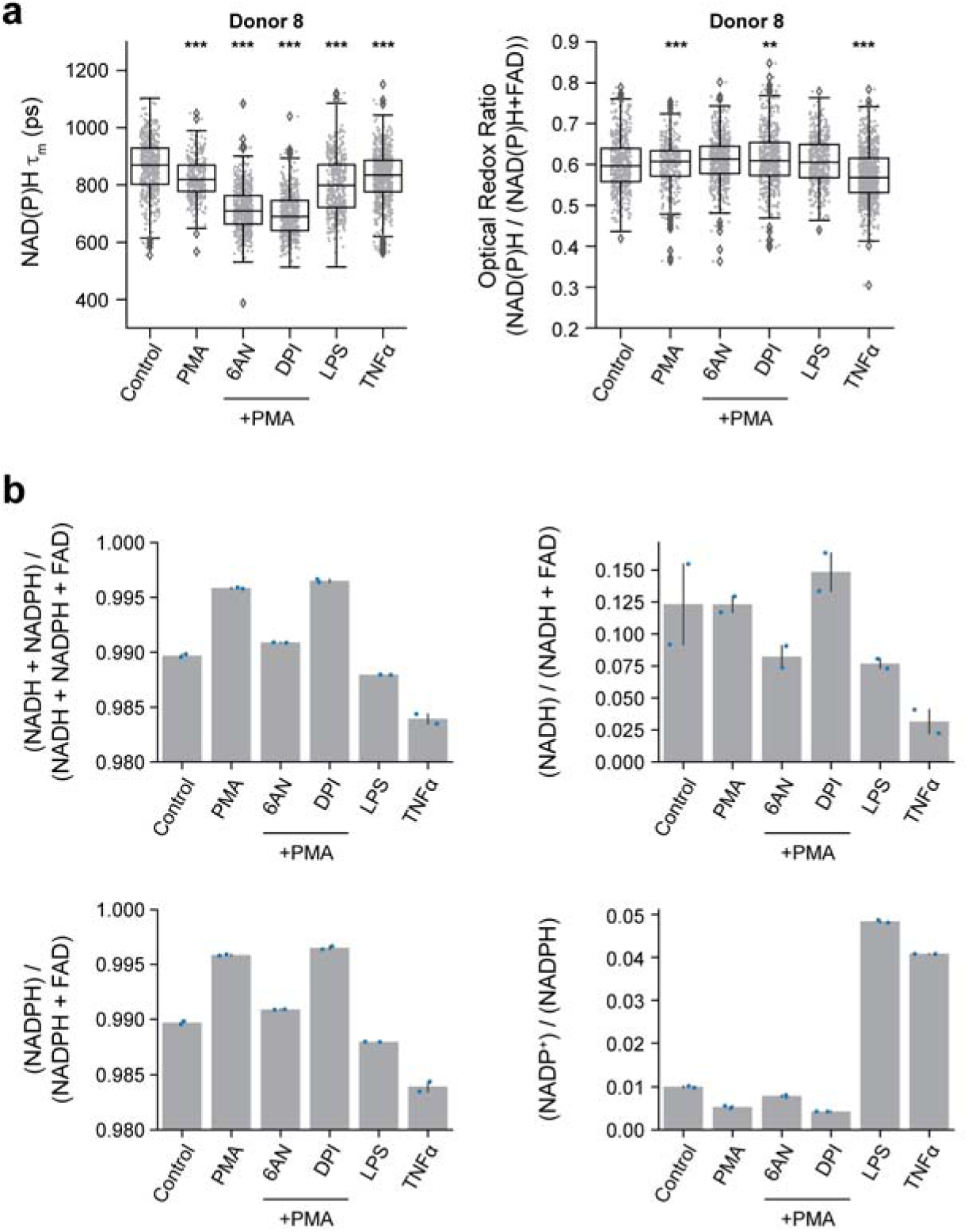
Paired OMI and LC-MS measurement of cells from the same donor (Donor 8). (a) Single cell quantification of NAD(P)H mean lifetime (left) and optical redox ratio (right) of control and PMA (100nM) and PMA plus inhibitor (5mM 6AN or 10µM DPI), LPS (20µg/L) and TNFα (5µg/L) treated neutrophils where each point is a single cell. Statistical significance of differences between multiple conditions for data presented in (a) were tested using ANOVA with posthoc Tukey’s test (*** p< 0.001; ** p< 0.01; * p< 0.05). Error bars represent the 95% confidence interval. (b) Redox ratios computed from LCMS measurements for the indicated conditions for two technical replicates. All data were collected 15 minutes after treatment.

**Supplementary Figure 8.**
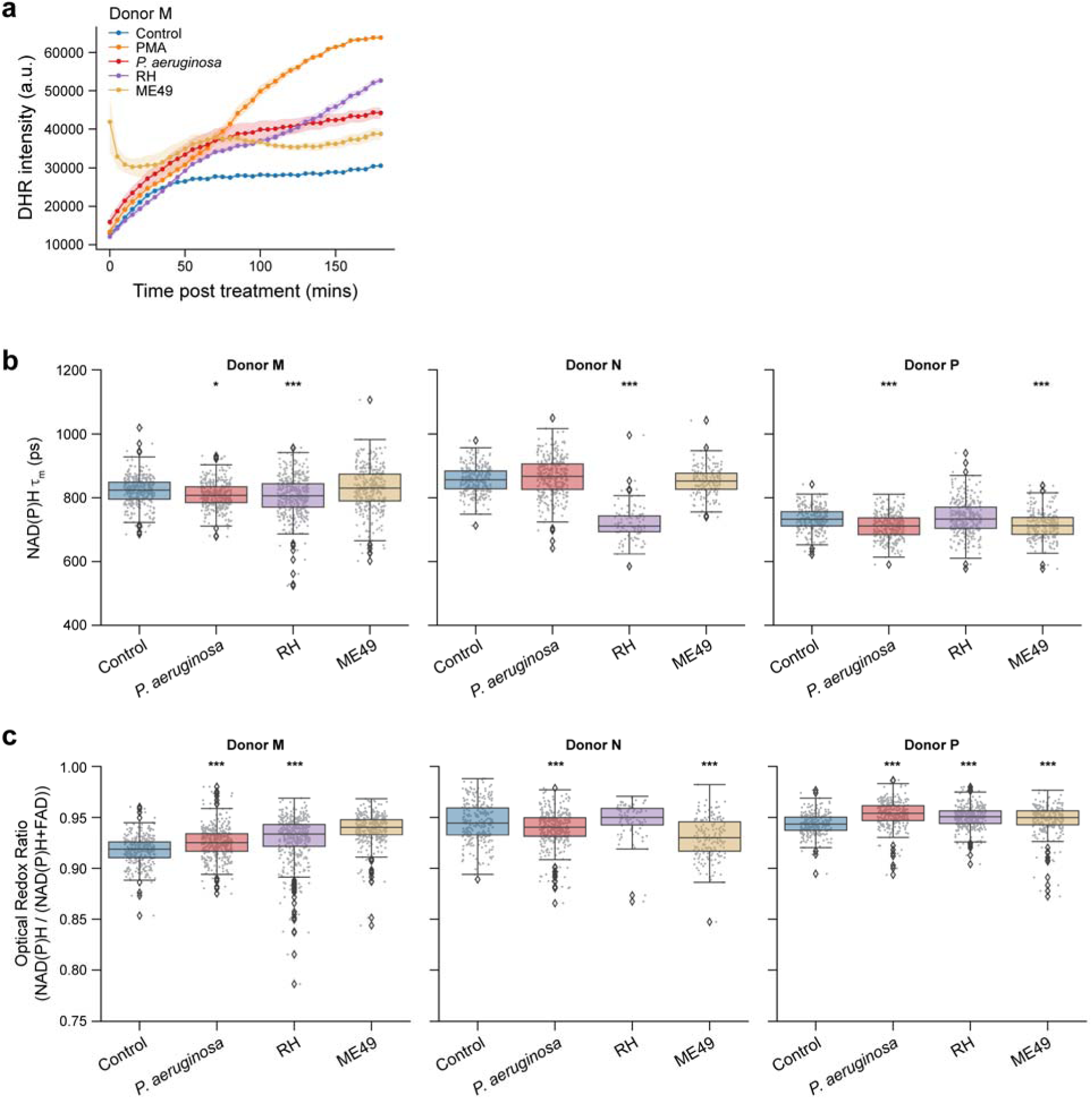
(a) Quantification of fluorescence intensity of Dihydrorhodamine 123 (DHR) indicating intracellular ROS in unstimulated control, PMA, coculture with *P. aeruginosa*, and *T. gondii* strains RH and M49. This is the 2nd repeat of the experiment with a different donor (Donor M) compared to data presented in Fig. 4a. (b) NAD(P)H mean lifetime and (c) optical redox ratio separately plotted for the 3 distinct donors (Donor M, N and P) for indicated conditions. At least 5 images were acquired per condition and each data point is a single cell. Number of cells in Table 1. Significance is tested using ANOVA with posthoc Tukey’s test (*** p< 0.001; ** p< 0.01; * p< 0.05). Data presented in Figs. 4b and 4c are derived from (b) and (c) respectively. Data in (b-c) were collected 15 minutes after treatment.

**Supplementary Figure 9.**
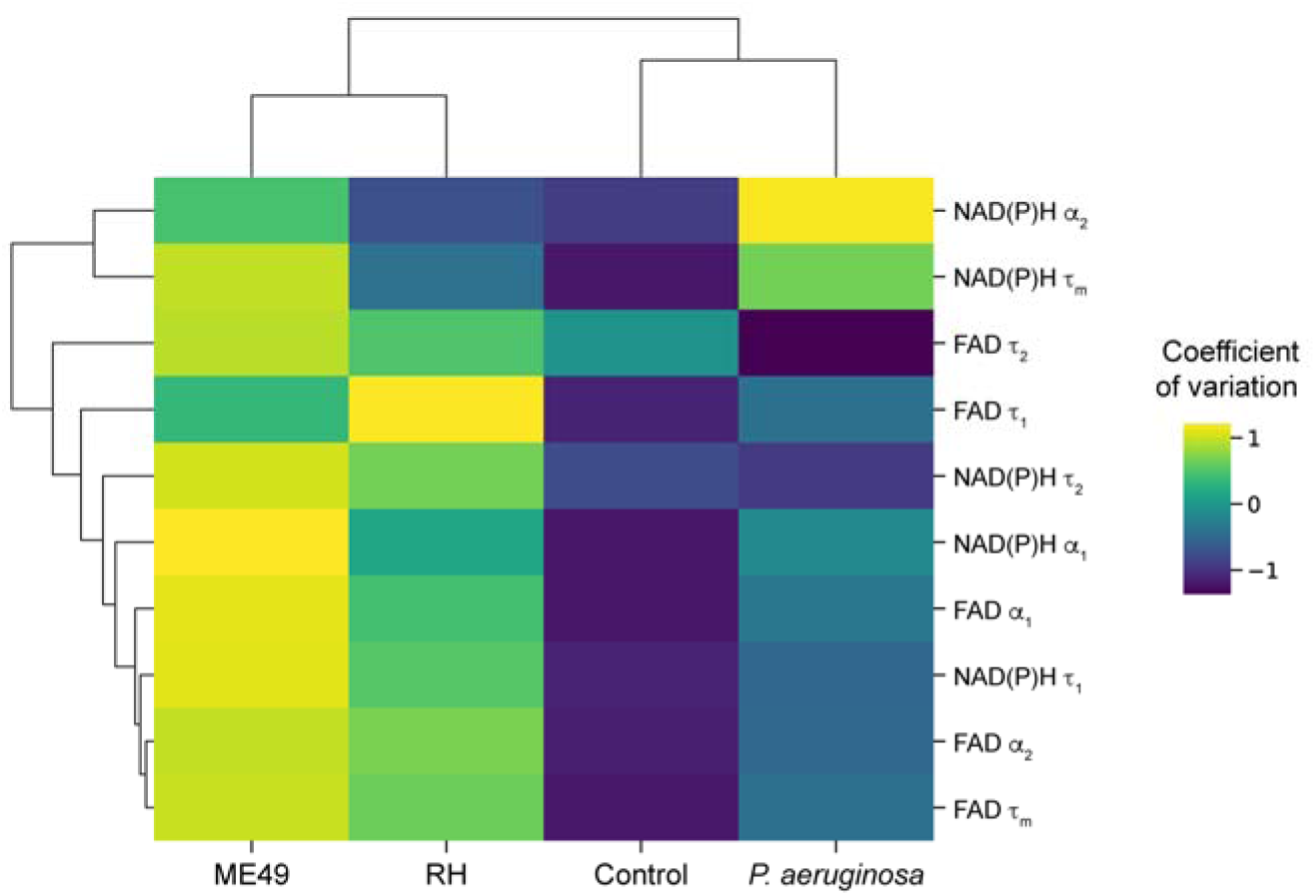
Heatmap showing coefficient of variation of OMI variables (subset of variables presented in Fig. 4e) in unstimulated control, and neutrophils activated with *P. aeruginosa*, and *T. gondii* strains RH and ME49. All plots include data from 3 distinct donors (Donor M, N and P, number of cells per donor in Table 1). All data collected at 15 minutes after infection.

